# Zfp503/Nlz2 is Required for RPE Differentiation and Optic Fissure Closure

**DOI:** 10.1101/2022.03.28.486078

**Authors:** Elangovan Boobalan, Amy H. Thompson, Ramakrishna P. Alur, David McGaughey, Lijin Dong, Grace Shih, Emile R. Vieta-Ferrer, Ighovie F. Onojafe, Vijay K. Kalaskar, Gavin Arno, Andrew J. Lotery, Bin Guan, Chelsea Bender, Omar Memon, Lauren Brinster, Clement Soleilhavoup, Lia Panman, Tudor C. Badea, Andrea Minella, Antonio Jacobo Lopez, Sara Thomasy, Ala Moshiri, Genomics England Research Consortium, Delphine Blain, Robert B. Hufnagel, Tiziana Cogliati, Kapil Bharti, Brian P. Brooks

## Abstract

**Purpose:** Uveal coloboma is a congenital eye malformation caused by failure of the optic fissure to close in early human development. Despite significant progress in identifying genes whose regulation is important for executing this closure, mutations are detected in a minority of cases using known gene panels, implying additional genetic complexity. We have previously shown knock down of *znf503* (the ortholog of mouse Zfp503) in zebrafish causes coloboma. Here we characterize *Zfp503* knock out (KO) mice and evaluate transcriptomic profiling of mutant vs. wild-type (WT) retinal pigment epithelium (RPE)/Choroid.

**Methods:** *Zfp503* KO mice were generated by gene targeting using homologous recombination. Embryos were characterized grossly and histologically. Patterns and level of developmentally relevant proteins/genes were examined with immunostaining/*in situ* hybridization. The transcriptomic profile of E11.5 KO RPE/choroid was compared to that of WT.

**Results:** *Zfp503* is dynamically expressed in developing mouse eyes and that loss of its expression results in uveal coloboma. KO embryos exhibit altered mRNA levels and expression patterns of several key transcription factors involved in eye development, including *Otx2, Mitf, Pax6, Pax2, Vax1* and *Vax2,* resulting in reduced melanin pigmentation in the presumptive RPE and its differentiation into neural-retina-like lineages. Comparison of RNA-Seq data from wild type and KO E11.5 embryos demonstrated reduced expression of melanin-related genes and significant overlap with genes known to be dynamically regulated at the optic fissure.

**Conclusions:** These results demonstrate a critical role of *Zfp503* in RPE differentiation and in optic fissure closure.

## Introduction

The optic fissure is a transient ventral opening along the optic cup (OC) and optic stalk (OS) during early ocular morphogenesis that permits the migration of periocular mesenchyme (Me) into the developing eye^1, 2^. Proper eye development requires that the outer edges of the optic fissure, composed of a single layer of presumptive retinal pigment epithelium (pRPE) cells, approximate and fuse during the fifth to seventh week of human gestation, equivalent to embryonic days (E)10.5-12.5 in the mouse^3^. Failure to do so results in uveal coloboma, a potentially blinding developmental eye defect in which a ventral portion of the iris, retina/RPE/choroid and/or optic nerve does not properly form^4^. Uveal coloboma may be isolated or part of a systemic syndrome such as CHARGE (<u>c</u>oloboma, <u>h</u>eart anomaly, choanal <u>a</u>tresia, <u>r</u>etardation, <u>g</u>enital and <u>e</u>ar anomalies)^5, 6^. Mutations in more than 20 different genes have been associated with uveal coloboma. These mutations, however, account for less than 10% of cases in the absence of a clear syndromic diagnosis^7, 8^, implying that additional genes and factors are involved.

We have reasoned that genes that are dynamically-regulated at the edges of the optic fissure during the process of optic fissure closure are likely to be important for the execution of this developmental process^9^. Such genes, we posit, are candidates for human coloboma. We identified 168 differentially regulated genes between E10.5 and E12.5 in wild type (WT, C57BL6/J) mice and demonstrated that morpholino knockdown in zebrafish of one such gene, *nlz2*, the ortholog of mouse *Zfp503*, results in coloboma through a *pax2*-dependent mechanism ^9, 10^. In mouse, *Zfp503* (*Nlz2, Zeppo2, Nolz1, Znf503)* mRNA is expressed throughout the OC at E10.5, becoming more limited to the area around the optic fissure by E11.5, and undetectable by *in situ* hybridization by E12.5 ^9^. Similarly, Nlz2 mRNA is expressed at the margins of optic fissure at 24 hpf and is no longer detectable at 48 hpf ^10^. Both *Zfp503* and another, closely-related zinc-finger motif gene, *Zfp703* (*Nlz1, Znf703, Zeppo1*) are homologous to the *Drosophila nocA* (*no ocelli*) gene^11^. The nomenclatures of both *Zfp503* and *Zfp703* are quite heterogeneous in the literature. Indeed, the paralogues of these genes in humans, *ZNF503* and *ZNF703,* respectively, do not follow the same conventions in databases such as Ensembl and the UCSC Genome Browser (accessed October 2021). To avoid confusion, we will refer to the mouse genes by the preferred convention *Zfp503, Zfp703* throughout the remainder of this paper. ZFP503 and ZFP703 proteins contain a single zinc finger, a Buttonhead domain, and an Sp-1-like domain, and act as transcriptional repressors in most studied systems^12–17^. Although first identified in zebrafish, *Zfp503/Znf503* is widely expressed during embryogenesis in the brain, spinal cord, face, limbs, and somites of zebrafish, mice, and chicken^12, 16–22^. During embryogenesis, *Zfp503/Znf503* is involved in a number of developmental processes, including motor neuron identity specification^12^, limb formation^21^, hindbrain patterning^19^, and striatal development^20, 23^. *Zfp503* is also expressed in adult mammary epithelial tissue, where it downregulates E-cadherin and GATA3, promoting an aggressive cancer phenotype^13, 14^.

In this study, we created a knockout (KO) mouse line (*Zfp503^-/-^*), resulting in complete loss of the corresponding protein, to gain further mechanistic insight into the role of *Zfp503* during eye development. *Zfp503^+/-^* mice were viable and exhibited congenital optic nerve excavation. Mouse KO embryos displayed near complete loss of melanin pigment in the pRPE, and uveal coloboma with 100% penetrance. Profiling of developmentally regulated ocular transcription factors revealed down-regulation of MITF and OTX2 and up-regulation and/or anatomically expanded expression of PAX6, PAX2, VSX2 proteins, and *Vax1* and *Vax2* transcripts, particularly in the ventral, proximal pRPE, accompanied by an expansion of cell number. These results were confirmed and quantified at the mRNA level in pRPE of E11.5 KO and WT embryos. *Zfp503^-/-^* mice were never observed in born litters, likely because of lung, and/or rib cage abnormalities. Gene expression profiling by RNA sequencing (RNA-Seq) revealed significant downregulation of melanin pigment-related genes and changes in multiple genes known to be dynamically regulated at the closing edges of the optic fissure. Sequencing of a cohort of patients with uveal coloboma did not reveal convincing loss-of-function alleles, consistent with lack of postnatal viability in *Zfp503^-/-^* mice.

## Materials and Methods

### Generation of Zfp503^-/-^ Mice

*Zfp503^-/-^* mice were generated by gene targeting using homologous recombination. The entire coding sequence of *Zfp503* (NM_145459.3) spanning exon 2 and exon 3 was replaced by a β-galactosidase-pGK-Neo cassette (Supplemental Figure S1A). The targeting vector was electroporated into W4 (SV129/S6) ES cells. ES clones containing the correctly targeted DNA sequence were identified by long genomic PCR and Southern blot. ES clones were further examined by karyotyping (data not shown) and injected into blastocysts of recipient mice.

Germline transmission was achieved as evidenced by approximately 90% F1 offspring exhibiting agouti coat color. Tail DNA genotyping of *Zfp503* WT allele was done with primers 5’-Exon2 ORF(GGCCACCCAGGATTATTC) and 3’-Exon2 ORF (GCCGCCGCCGCTGTGCTTAC) and mutant allele was genotyped with primers 5’-Exon2 ORF(GGCCACCCAGGATTATTC) and 3’-beta-galactosidase gene (CGGGCCTCTTCGCTATTACG). The WT band was 216 bp and the mutant band was 385 bp long. Heterozygous mice were backcrossed into C57BL/6J background for at least five generations. The floxed Neo cassette was removed by mating heterozygous *Zfp503* mice with *Zp3*-*Cre* mice.

### Animal Husbandry

Mice were housed and maintained in a 14 hr light, 10 hr dark cycle. All experiments were conducted in accordance with the Animal Study Protocol (ASP) NEI-605 approved by the NEI Animal Care and Use Committee as well as the ARVO Statement for the Use of Animals in Ophthalmic and Vision Research. Clinical examination of the anterior segment of the eyes was performed on gently restrained awake mice using a Haag-Streit BQ slit lamp (Bell Ophthalmic Technology, Westville, NJ, USA). After pupil dilation with one drop of 1% tropicamide (Alcon Laboratories, Inc., Fort Worth, TX, USA), the posterior segment of the mouse eyes was examined with a 90-D condensing lens (Volk, Mentor, OH, USA). Fundus images were obtained using the Micron III imaging system (Phoenix Research Laboratories, CA, USA) on mice sedated with intraperitoneally injected ketamine (100 mg/mL) and xylazine (200 mg/mL) diluted in saline. The eyes were dilated with 0.5% tropicamide (Alcon Laboratories, Fort Worth, TX, USA), 2.5% phenylephrine hydrochloride for local muscle relaxation and 0.5% proparacaine hydrochloride for local anesthesia. Hypromellose ophthalmic demulcent solution (Gonak) was used to prevent dehydration. After imaging, mice were kept on a heating pad until they were awake. Mice were euthanized with carbon dioxide using institutional standards. For histology, eyes were dissected, fixed overnight in 10% formalin and embedded in methacrylate for sectioning through the pupillary-optic nerve axis and staining with hematoxylin and eosin (H&E).

### Immunofluorescence

Embryos were washed in PBS for 5 min and fixed in 4% paraformaldehyde in PBS for 0.5-3 hr, then washed with PBS and infiltrated with 30% sucrose overnight at 4°C. Next, they were cryopreserved in Tissue Freezing Medium (Thermo Fisher Scientific, Baltimore, MD) and 16 µm cryosections were prepared with a Leica Biosystems CM3050 S cryostat (Buffalo Grove, IL). Assays were repeated multiple times as indicated in the text and Figure Legends on at least three independent cryosections. Indirect immunofluorescence was performed with the following antibodies: rabbit polyclonal anti-ZNF503 (Sigma, St. Louis, MO, HPA026848, 1:1000), rabbit polyclonal anti-Pax6 (Covance, Princeton, NJ, 1:1000), rabbit polyclonal anti-Pax2 (Invitrogen, Carlsbad, CA, 1:500), goat polyclonal anti-OTX2 (R&D Systems, Minneapolis, MN, AF1979, 1:2000), goat polyclonal anti-Chx10/Vsx2 (Santa Cruz Biotechnology, Inc., Dallas, Texas 75220, SC-21690, 1:2000), and rabbit anti-Mitf (gift from K. Bharti) ^24^. For all the antibodies used in this study, antigen unmasking was achieved by boiling the tissue sections in target retrieval solution (Agilent/Dako S1700, Santa Clara, CA) for 20 min prior to incubation with the primary antibody. Following overnight incubation in primary antibody, sections were washed three times for 10 min with phosphate buffered saline with 0.1% Tween (PBST, Invitrogen, Camarillo, CA.) and incubated with Alexa-Flour-488 or 555 secondary antibodies (1:1000) along with 4’,6-diamidino-2-phenylindole (DAPI, Invitrogen, Camarillo, CA, 1:1000) for 1 hr at room temperature (RT). Next, sections were washed with PBST (3 times for 10 min) and mounted with a coverslip. Images were acquired with a Zeiss LSM700-880 confocal microscopes (Carl Zeiss Microscopy, LLC, Thornwood, NY) maintaining the same settings and parameters when comparing two or more groups. Details of primary antibodies used are also provided in Supplemental Table S1.

### In Situ Hybridization

RNA *in situ* hybridization was performed using the RNAScope® Assay, Multiplex Fluorescent Reagent Kit V2 (Advanced Cell Diagnostics, ACD, Newark, CA). Briefly, fixed embryo cryosections were dehydrated with graded alcohols (50%, 70% and 100%) for 5 min each followed by hydrogen peroxide treatment (Cat. No. 322381, ACD, Newark, CA) for 10 min at RT. The slides were then washed with distilled water and antigen retrieved with RNAScope® 1x Target Retrieval Reagent in a steamer for 5 min. After washing with distilled water, the slides were dehydrated with 100% ethanol. The samples were then treated with RNAscope® Protease III for 30 min at 40°C and washed with distilled water. Mouse *Vax1* (Cat. No. 85101-C2) and *Vax2* (Cat. No. 535091) RNAscope® Target probes were mixed at a ratio of 1:50 and hybridized for 2 hr at 40°C followed by two 2-min wash with 1x wash buffer (Cat. No. 310091, ACD). For signal amplification, sections were incubated with Amp1 for 30 min at 40°C and washed twice for 2 min with 1x wash buffer, followed by similar treatment with Amp2 and Amp3 for 15 min each and washes. In the last two-step of amplification, the sections were treated first with HRP-C1 for 15 min at 40°C, washed twice for 2 min with 1x wash buffer and incubated with Tyramide Signal Amplification (TSA^TM^) plus fluorescein (1:1500) for 30 min at 40°C. After rinsing twice with 1x wash buffer and blocking with HRP-blocker for 15 min at 40°C followed by two rinses with 1x wash buffer, the slides were similarly treated with TSA^TM^ plus Cyanine 3 (1:1500). Nuclei were visualized with DAPI (Cat. No. 310098, ACD, Newark, CA) for 30 seconds at RT. Slides were mounted with Prolong Gold antifade mounting medium and imaged with a Zeiss LSM800 Confocal microscope (Carl Zeiss Microscopy, LLC, Thornwood, NY) maintaining the same settings and parameters when comparing two or more groups.

### Beta-galactosidase Staining

E11.5 mouse embryos were fixed with 0.2% glutaraldehyde overnight at 4°C, followed by 3x15 min wash with Buffer A (100 mM PBS, 2 mM MgCl2, 5 mM EGTA) at RT and 2x10 min wash with Buffer B (100 mM PBS, 2 mM MgCl2, 0.01% sodium deoxycholate, 0.02% NP40). Color was developed in the dark by incubating the embryos with Buffer C (5 mM potassium ferricyanide, 5 mM potassium ferrocyanide, 1 mM x-Gal) at 37°C. The embryos were rinsed with PBS and imaged with a Leica M205 FA fluorescence stereo microscope, (Leica Microsystems Inc., Buffalo Grove, IL, USA).

### Skeletal Staining

Mouse embryos at E18.5 were stained with Alcian Blue 8GX (A0298, Bio Basic Canada, Inc. Markham ON, Canada) and Alizarin Red S (SC-205998, Santa Cruz Biotechnology, Inc. Dallas, Texas 75220, USA) as described previously ^25^. Embryos were skinned and eviscerated after fixation in 95% ethanol overnight. Three complete E18.5 skeletons were stained for each WT and *Zfp503^-/-^* samples. The arms and legs of the skeletons were removed to facilitate the visualization of the sternum. Photographs were acquired with a Leica MC190 HD digital camera.

### ZFP503 Human DNA Sequencing

Uveal coloboma patients and, when available, their first-degree relatives, were recruited and examined in the Ophthalmic Genetics Clinic of the National Eye Institute under an IRB-approved protocol (NCT00368004, NCT01778543 or NCT00076271, www.clinicaltrials.gov). DNA from either peripheral blood leukocytes or saliva was screened for pathogenic variants in genes associated with uveal coloboma, using previously described methods and workflow ^26^. Briefly, the NEXTFLEX Rapid XP DNA-Seq kit (PerkinElmer) and xGen Lockdown probes (Integrated DNA Technologies, Inc, Coralville, Iowa, USA) were used to capture exons and other genomic regions with known or suspected pathogenic variants from a custom panel of 731 genes implicated in eye development or disease. The libraries were then sequenced on Illumina NextSeq￼550(Illumina￼San Diego, CA, USA), aligned, variants (including copy number variations) called, annotated, and prioritized through a custom pipeline available on GitHub. The pathogenicity and effect of variants were interpreted according to the ACMG guidelines ^27^. For the independent, phenotypically agnostic cohort, the affected individual was consented and underwent whole genome sequencing as part of the United Kingdom 100,000 Genomes Project (UK100KGP) as previously described ^28^. Both parental samples were not available for segregation analysis. The subject’s sequence underwent variant interpretation using the retinal disorders virtual gene panel (version 2.21, https://panelapp.genomicsengland.co.uk/panels/307/) which failed to identify a causative genotype in any known inherited retinal degeneration genes. Loss of function variants in the *ZNF503* gene were interrogated in the UK1000KGP Interactive Variant Analysis (v2.0) dataset including 45,356 individual genomes. High confidence loss of function variants (frameshift, nonsense and canonical splice variants) with a minor allele frequency <0.001 in the gnomAS dataset were identified in the genomes and confirmed by visualization on the individual paired-end reads using the Integrative Genomics Viewer ^29^. https://github.com/Bin-Guan). The pathogenicity and effect of variants were interpreted according to the ACMG guidelines ^27^. For the independent, phenotypically agnostic cohort, the affected individual was consented and underwent whole genome sequencing as part of the United Kingdom 100,000 Genomes Project (UK100KGP) as previously described ^30^. Both parental samples were not available for segregation analysis. The subject’s sequence underwent variant interpretation using the retinal disorders virtual gene panel (version 2.21, https://panelapp.genomicsengland.co.uk/panels/307/) which failed to identify a causative genotype in any known inherited retinal degeneration genes. Loss of function variants in the *ZNF503* gene were interrogated in the UK1000KGP Interactive Variant Analysis (v2.0) dataset including 45,356 individual genomes. High confidence loss of function variants (frameshift, nonsense and canonical splice variants) with a minor allele frequency <0.001 in the gnomAS dataset were identified in the genomes and confirmed by visualization on the individual paired-end reads using the Integrative Genomics Viewer ^29^.

### RNA-Sequencing

Embryos (E11.5) were harvested after euthanizing timed pregnant Zfp503^+/-^ mice with CO_2_. Eyes were enucleated and the RPE/choroidal tissues were dissected after removing the retina and lens. The RPE/choroidal tissues were stored in RNAlater (Qiagen, Germantown, MD, USA) at 4°C. Three samples per genotype (each pooled from four embryos) were lysed and total RNA was isolated using RNAeasy mini kit (Qiagen). RNA integrity was verified on a Bioanalyzer (Agilent Technologies, Santa Clara, CA, USA) and samples with RIN 8 and greater were processed for RNA-Seq by the NIH Intramural Sequencing Center (NISC).

STAR 2.4.2a was used with GRCm38 to align the RNA-Seq data (https://pubmed.ncbi.nlm.nih.gov/23104886/). Counts were quantified with HTSeq 0.6.1p1 with the gencode.vM8.primary_assembly.annotation.gtf (https://pubmed.ncbi.nlm.nih.gov/25260700/). DESeq2 1.30.0 was used for RNA-Seq differential analysis (https://pubmed.ncbi.nlm.nih.gov/25516281/). As predicted, relatedness of gene expression was greater within either *Zfp503^+/+^* or *Zfp503^-/-^* samples than between the two groups (Supplemental Figure S2). PCA analysis was performed using the base R (4.0.5) prcomp function with the rlogTransformation function and the 500 most variable genes. This analysis demonstrated notable differences between the *Zfp503^-/-^* and *Zfp503^+/+^* groups, with PC1 explaining 80% of the total variance (data not shown). For differential testing, the design was set as “∼condition + PCA2 + laneID” where condition is the genotype status of the mouse (e.g., *Zfp503^-/-^* or *Zfp503^+/+^*) and PCA2 and sequencing laneID were covariates to be corrected for.PCA2 was used as a covariate as the separation of the samples was not related to any biological factor. The DESeq2 nbinomWaldTest was used to calculate the *p* values (https://pubmed.ncbi.nlm.nih.gov/25516281/) and the corrected *p* values were calculated with the fdrtool function (https://pubmed.ncbi.nlm.nih.gov/18441000/).

### Data availability

RNA-Seq data have been deposited under GEO accession number GSE180641: https://www.ncbi.nlm.nih.gov/geo/query/acc.cgi?acc=GSE180641.

## Results

### Zfp503 is expressed in the developing mouse eye

We have previously shown that *Zfp503* mRNA is expressed throughout the optic vesicle and subsequently, it is confined to the fusing optic fissure, before becoming undetectable by *in situ* hybridization in the developing zebrafish and mouse eye ^9^. To confirm the pattern of protein expression in the developing mouse, we performed immunofluorescent labeling of coronal cryosections of WT embryos at E10.5 (open fissure), E11.5 (closing fissure) and E12.5 (closed fissure) (Figure 1). ZFP503 expression was prominently seen in the pRPE and surface ectoderm, with qualitatively lower levels of expression in the distal optic stalk at E10.5. From E11.5 through E12.5, the pRPE and the optic stalk both showed prominent ZFP503 expression.

**Figure 1.**
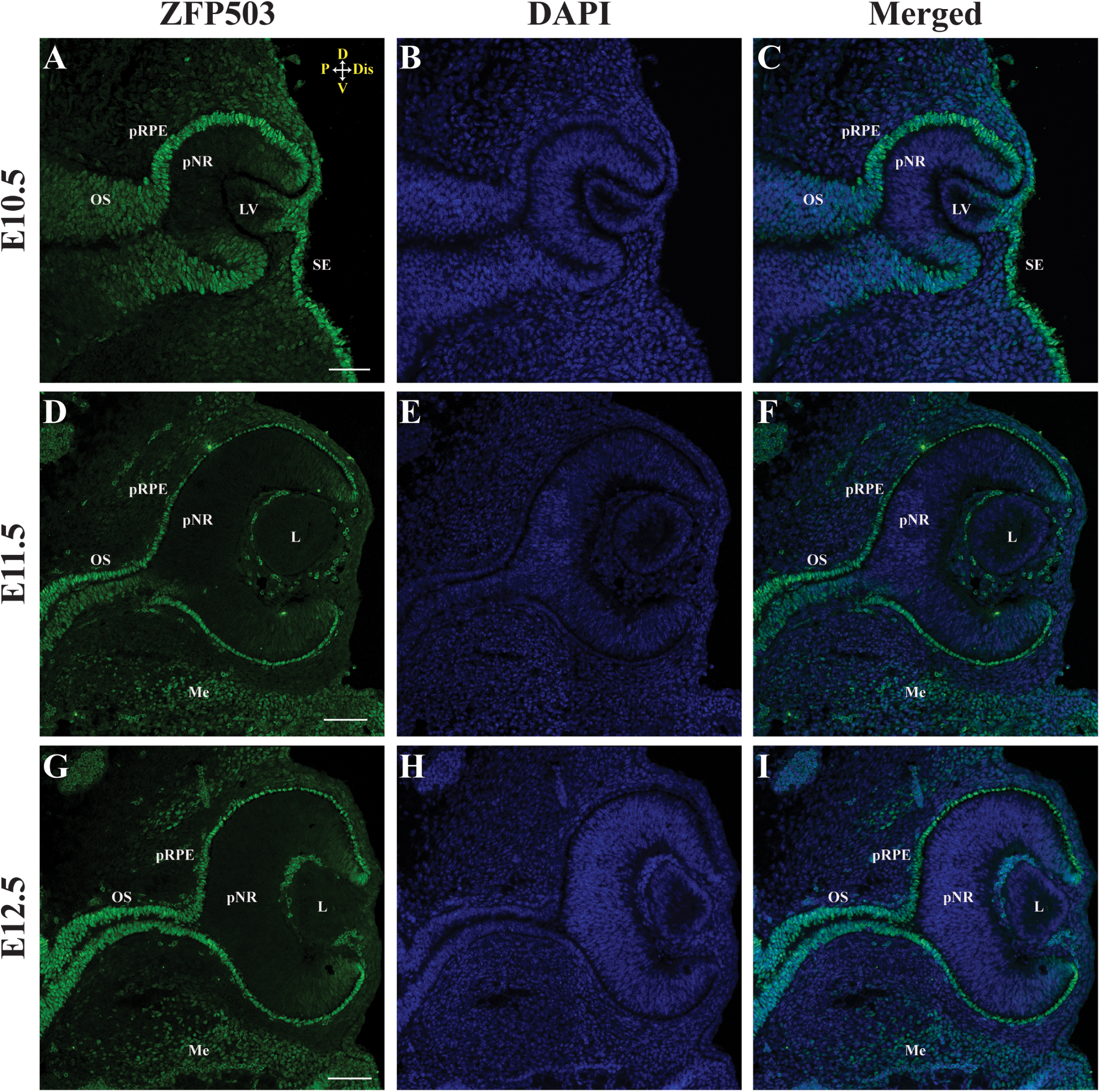
ZFP503 protein is expressed in the developing mouse eye. ZFP503 expression (green) by immunofluorescence in representative coronal sections of WT mouse embryos at three developmental time points corresponding to “before closure” at embryonic day (E)10.5, “during closure” (E11.5) and “after closure” (E12.5) of the optic fissure (n=3 embryos each). DAPI nuclear staining in blue. Compass in (A) applies to all panels: D, dorsal; V, ventral; P, proximal; Dis, distal. OS, optic stalk; pRPE, presumptive retinal pigment epithelium; pNR, presumptive neural retina; LV, lens vesicle; L, lens; Me, Mesenchyme; SE, surface ectoderm. Scale bar, 100 μm.

Sagittal sections of the OC (Supplemental Figure S3) revealed that ZFP503 was expressed in the pRPE lining the optic fissure at E11.5 and that, by E12.5, a small patch of faint expression remained in the presumptive neural retina (pNR) at the now fused margins. Variable levels of expression were observed in the local Me, especially ventral to the OC (Figure 1). We describe Me expression as “local” and not “periocular”, as there was generally a layer of Me that did not express ZFP503 immediately adjacent to the OC. In mature retina, ZFP503 expression was detectable in a sub-population of cells in the ganglion cell layer and in cells of the inner nuclear layer, presumably a subset of amacrine cells (Supplemental Figure S4). Notably, the labelled cells did not uniquely coincide with a number of known ganglion cell and amacrine cell subpopulations i.e., BRN3a-positive (Supplemental Figure S4A-C, A’-C’, arrows pointing to double-labeled cells), ISL1-positive (Supplemental Figure S4D-F, D’-F’, arrows pointing to double-labeled cells), and CALRETININ-positive (Supplemental Figure S4G-I, G’-I’, arrow pointing to a single double-labeled cell).

### Targeted deletion of Zfp503 results in systemic abnormalities

To better assess the role of *Zfp503* in mouse eye development, we established the *Zfp503^-/-^* mouse line via targeted deletion of the *Zfp503* coding sequence contained in Exon2 and Exon3 (Supplemental Figure S1A). Long-range PCR was used to confirm the presence of the recombined allele in *Zfp503^+/-^* and *Zfp503^-/-^* compared to *Zfp503^+/+^* mice (Supplemental Figure S1B). Supplemental Figure S1C shows the corresponding PCR genotyping of tail DNA. Exon2 and Exon3 deletion were further confirmed by RNA-Seq (data not shown.) Confocal photomicrographs of E12.5 coronal sections immunostained with anti-ZFP503 antibody showed expression in the pRPE, optic stalk and Me in the developing eye of WT embryos (Supplemental Figure S1D) but undetectable levels of ZFP503 in *Zfp503**^-/-^*** littermates (Supplemental Figure S1E).

To establish overall *Zfp503* expression, whole E11.5 mouse embryos were stained for β-galactosidase that is expressed in *Zfp503^-/-^* mice with the same spatio-temporal pattern of the replaced *Zfp503* gene. Consistent with previous reports^18, 21–23^, we observed β-galactosidase activity in the spinal cord, midbrain, eye, and limited areas of the forebrain and hindbrain of *Zfp503^-/-^* but not *Zfp503^+/+^* E11.5 embryos (Figure 2A, B). No qualitative difference in the distribution and intensity of β-galactosidase activity was noted in heterozygous vs. homozygous embryos (data not shown).

**Figure 2.**
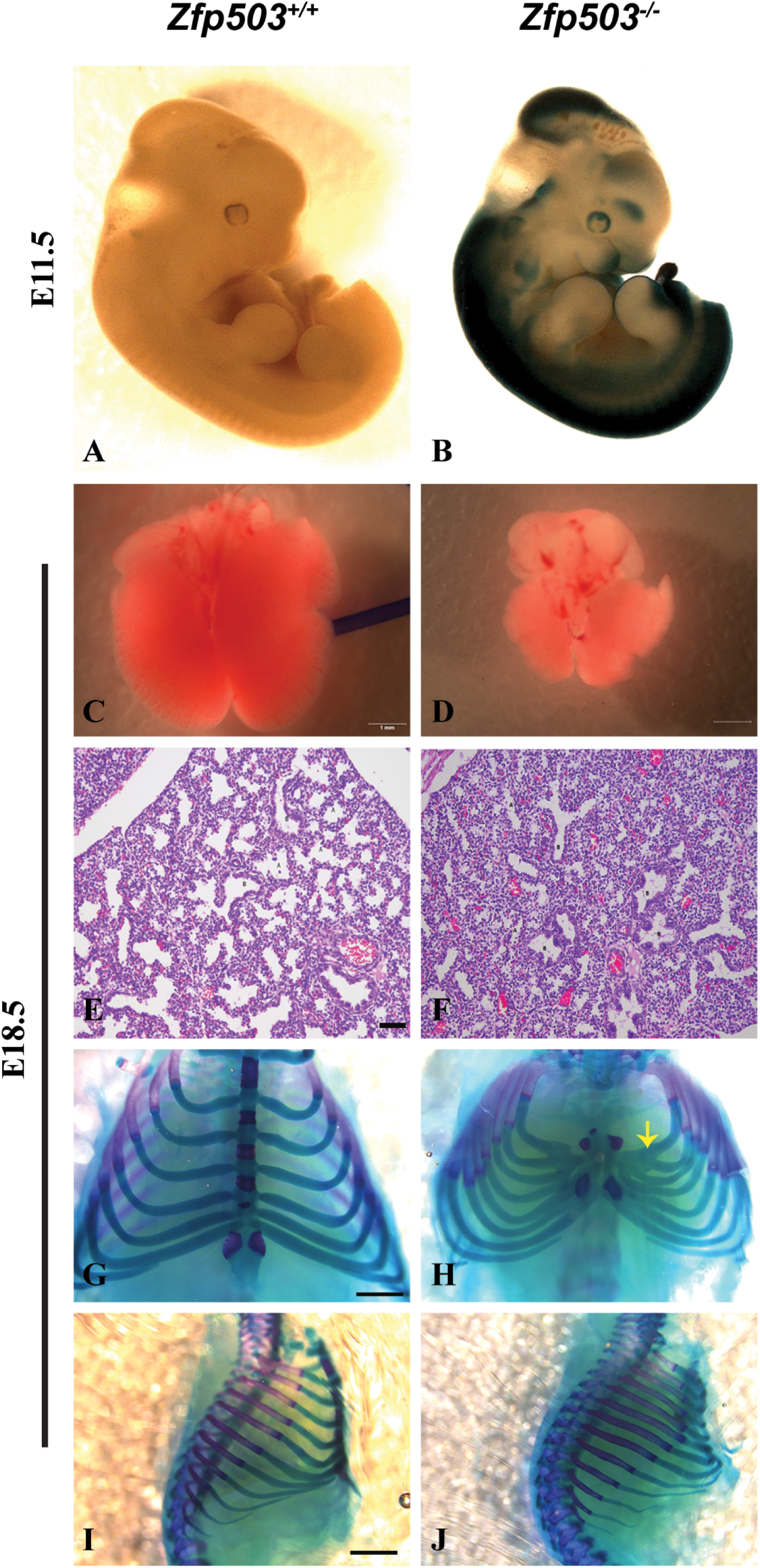
Homozygous deletion of *Zfp503* results in relatively normal appearing E18.5 embryos with characteristic organ pathologies. β-galactosidase staining (indicative of *Zfp503* localization) in representative *Zfp503^-/-^* embryos, was present in the developing brain, spinal cord, eye and branchial arches (A, B, n=4/genotype). Loss of ZFP503 resulted in smaller lungs (C, D, n=4/4 from two litters) and faulty alveolar formation in the lung (E, F, n= 3 sections through each of 4 embryos) visualized by H&E staining in representative histology sections. The sternum of *Zfp503^-/-^* embryos visualized with Alcian Blue was shorter than in WT embryos, accompanied by abnormal rib morphology (G, H frontal view, arrow, n=3/genotype) and (I, J lateral view). Scale bars, 1 mm (C and D); 50 μm (E and F); and 0.9 mm (G-J).

*Zfp503^+/-^* mice were viable, fertile and without obvious systemic abnormalities or dysmorphic features (data not shown). Although their anterior segment appeared normal, dilated fundus exam and ocular histology showed abnormal pigmentation and excavation surrounding the optic nerve (Supplemental Figure S5). Retinal layers, including the photoreceptor and RPE, were histologically normal, albeit some abnormal layering of pigment near the optic nerve head was noted. No major dysmorphic abnormalities were visible in *Zfp503^-/-^* embryos. However, homozygous deletion of *Zpf503* resulted in neonatal lethality. Viable KO embryos were observed as late as E18.5 (n=39) but were never observed at the time of weaning (n=555). This observation coupled with the broad expression pattern of *Zfp503* during development led us to inspect several major organ systems. Histology of heart, liver and brain were unremarkable in *Zfp503^-/-^* embryos (data not shown.) The lungs of E18.5 embryos of homozygous were notably smaller (n=4/4) compared to those of heterozygous/WT littermates (n=12) (Figure 2 C, D). Histologic examination of E18.5 *Zfp503^-/-^* mice (n=4/4) revealed atelectatic lungs, with reduced quantities of alveoli, and bronchioles that were closer together than those of WT (n=4) mice (Figure 2E, F). The sternum was much shorter in *Zfp503^-/-^* than in WT embryos (Figure 2G-J), with some fusion of the attached ribs (arrow). No obvious changes in rib identity, vertebrae and limb patterning were observed in any of the mice analyzed (data not shown).

### Decreased pigmentation, incomplete optic fissure closure, and RPE abnormalities in null mouse eyes

Examination under the dissecting microscope showed decreased, but not completely absent, pigmentation in the pRPE of *Zfp503^-/-^* eyes compared to WT, beginning at approximately E11.5 (Figure 3A, B, arrow). Additionally, while the optic fissure was fully fused at E13.5 in WT embryos (Figure 3C), *Zfp503^-/-^* embryos had incomplete fusion with an opening in the ventral side of the OC (Figure 3D, arrow). A similar phenotype was noted in *Zfp503^-/-^* embryos as late as E18.5 (data not shown), arguing against developmental delay being the cause of this finding. H&E-stained sagittal sections of WT embryos at E13.5 revealed a pigmented, smooth and continuous pRPE, comprised of a single layer of cells clearly delineated from surrounding tissue (Figure 3E). The location of the optic fissure was no longer apparent. In *Zfp503*^-/-^ sagittal sections, the pRPE lacked pigmentation throughout and multiple layers of cells resembling pNR were observed near the ventral optic fissure (Figure 3F, arrows). Furthermore, the optic fissure failed to close, resulting in a thin, but clear, division between the temporal and nasal halves of the OC (arrowhead). Collagen IV staining of the basement membrane confirmed that fissure closure occurred in E13.5 WT embryos (Figure 3G), but that separation remained in *Zfp503^-/-^* mutants (Figure 3H). Consistent with these observations, coronal sections of embryos at E14.5 stained with H&E also showed marked absence of pRPE pigmentation in *Zfp503^-/-^* compared to WT embryos (Figure 4A, B, arrows). A single layer of RPE cells was visible in WT embryo coronal (Figure 4A and C) and sagittal sections (Figure 4E). Hyperplasia in the pRPE was visible both from coronal and sagittal sections of *Zfp503^-/-^* embryos (Figure 4 B, D and F, arrows) and was mostly proximal, between the optic stalk and developing lens (Figure 4B, D). Furthermore, expression of the pro-retinogenic transcription factor VSX2/CHX10 was prominent in pRPE region at E12.5 (Figure 4D) and E14.5 (Figure 4F).

**Figure 3.**
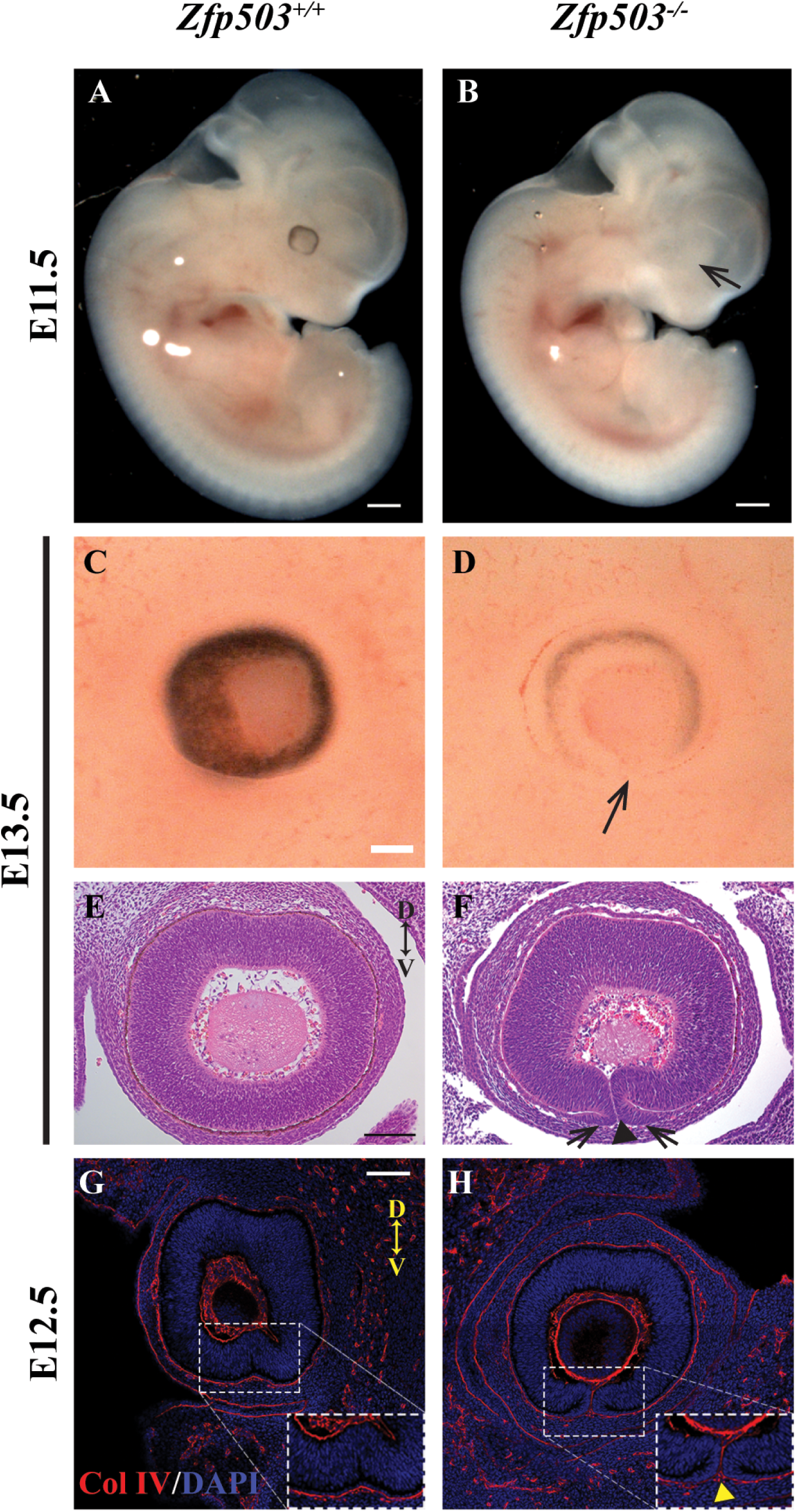
*ZFP503* deletion results in hypopigmentation of the RPE and uveal coloboma. Microphotographs of representative WT embryos at embryonic day (E)11.5 (A) and E13.5 (C) showed normal, circumferential and continuous pigmentation of the RPE that was almost completely absent in age-matched *Zfp503*^-/-^ embryos (B, D, arrows, n=5/genotype). Sagittal sections of representative E13.5 embryos stained with H&E revealed that the edges of the optic fissure were fused and no longer discernible in WT embryos (E), whereas representative *Zfp503^-/-^* embryos showed distinct non-fusion of fissure edges (F, arrowhead). Note the hyperplasia, resembling presumptive neural retina (pNR), flanking the fissure (arrows). Whereas the basement membrane of the optic fissure edges was no longer detectable by collagen IV immunostaining (red) in E13.5 WT embryos (G), a clear separation of the optic fissure edges could be seen in *Zfp503^-/-^* littermates (H, arrowhead, n=2/genotype). DAPI nuclear staining in blue. Confocal microscope exposure and parameters were maintained equal for comparison purposes. Compass in (E, G) applies to E-H: D, dorsal; V, ventral. Scale bars, 500 μm (A, B); 100 μm (C-H).

**Figure 4.**
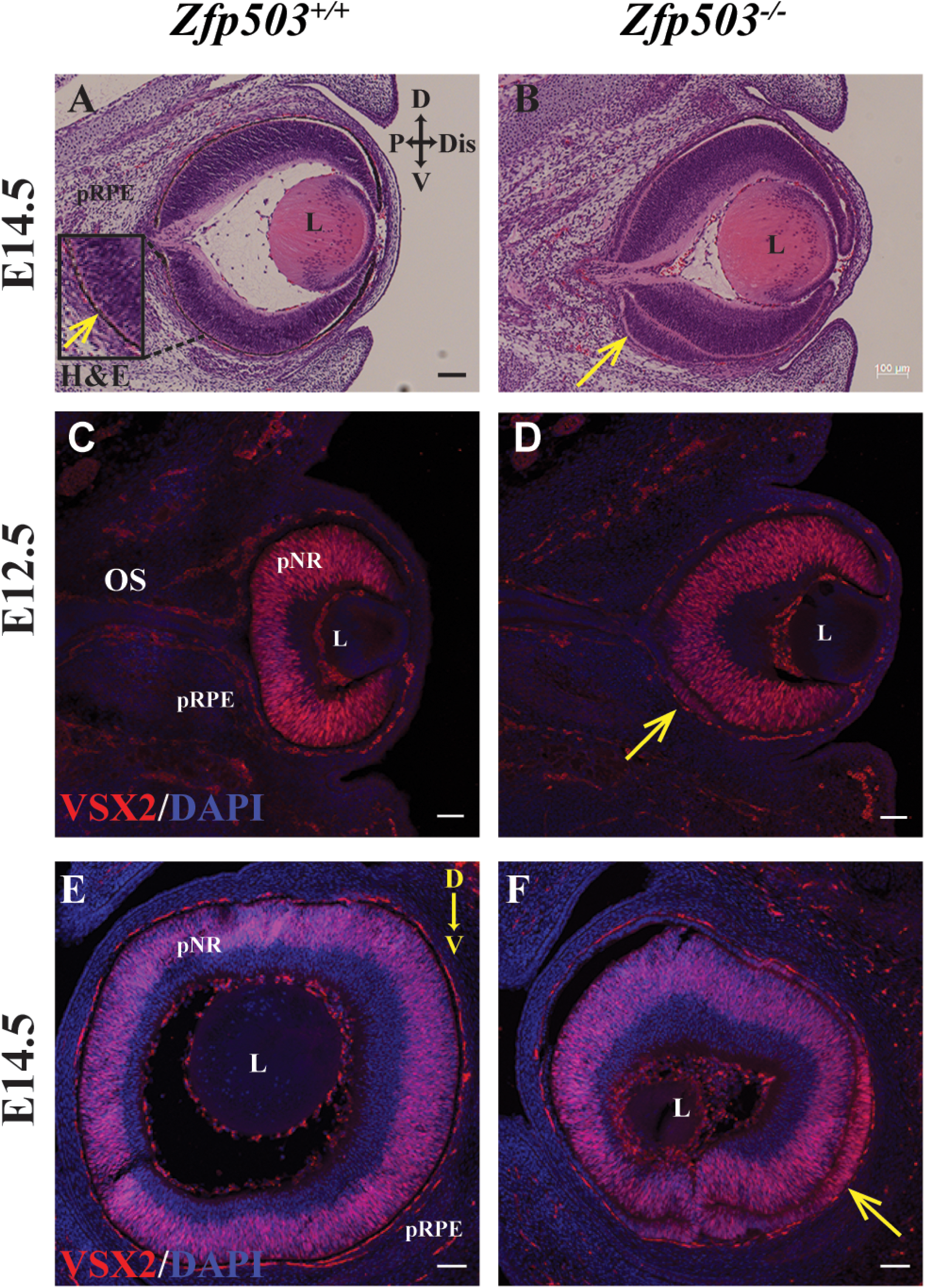
Hyperplasia is observed in the presumptive RPE (pRPE) of *Zfp503^-/-^* developing eyes. Representative coronal sections of embryonic day (E)14.5 WT (A) and *Zfp503^-/-^* embryos (B) stained with H&E displayed continued pRPE hypopigmentation with hyperplasia ventrally and proximally in *Zfp503^-/-^* (arrow, n=3/genotype). Upon fluorescent immunostaining (red), the pro-retinogenic transcription factor VSX2/CHX10 was detected in the neural retina at E12.5 (C, D, coronal sections, n=2/genotype) and E14.5 (E, F, sagittal sections, n=2/genotype), and abnormally in the hyperplastic pRPE in *Zfp503^-/-^* eyes (D, F, arrows). DAPI nuclear staining in blue. Confocal microscope exposure and parameters were maintained equal for comparison purposes. Compass in (A) applies to A-D, in (E) applies to E-F: D, dorsal; V, ventral; P, proximal, Dis, distal. pRPE, presumptive retinal pigment epithelium; OS, optic stalk; L, lens; pNR, presumptive neural retina. Scale bars,100 μm (A-D); and 200 μm (E, F).

### Targeted deletion of *ZFP503* results in abnormal patterns of transcription factor expression during eye development

Differentiation and maintenance of RPE involves a complex interplay of transcription factors and signaling molecules, including MITF, OTX2, VSX2, PAX6, VAX1, VAX2, WNTs, FGFs, and JAGGED-NOTCH signaling ^31–37^. Therefore, to investigate the mechanism by which *Zfp503^-/-^* ocular phenotypes occur, we performed immunostaining for several key developmental markers on cryosections of WT and *Zfp503^-/-^* embryos. pRPE is frequently identified molecularly by early expression of the transcription factors MITF and OTX2^38–41^. In WT embryos, MITF was expressed uniformly along the proximo-distal and dorso-ventral axes in the pRPE at E11.5 (Figure 5A, C, E). By contrast, MITF immunofluorescent signal was faint in the proximal OC in *Zfp503^-/-^* embryos (Figure 5B). Expression of MITF in KO embryos appeared to increase more distally yet remained relatively reduced compared to WT in the ventral-most portion of the pRPE near the optic fissure (Figure 5D, F). Of note, although MITF-positive cells normally line the optic fissure, MITF expression was absent to weak in *Zfp503^-/-^* eyes.

**Figure 5.**
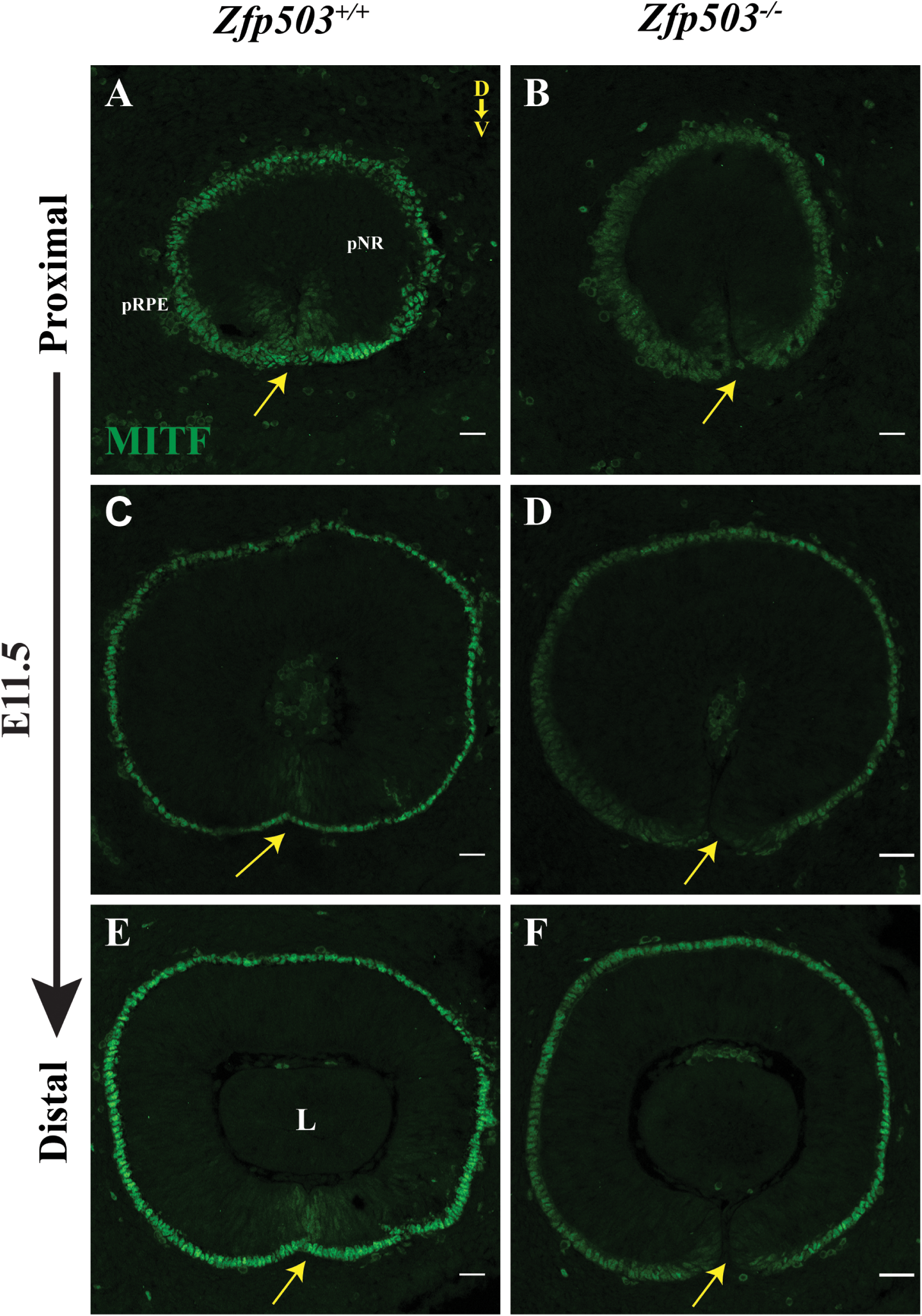
Loss of *Zfp503* is associated with reduced expression of MITF. Sagittal sections from representative embryonic day (E)11.5 WT (A, C, E) and *Zfp503^-/-^* (B, D, F) embryos immunostained with anti-MITF antibody (green) are shown in proximal to distal series. Note the open optic fissure (arrow) in all *Zfp503^-/-^* sections (B,D,F). Qualitative reduction of MITF expression in *Zfp503^-/-^* was particularly evident in proximal sections, near the optic stalk (B, arrow). All panels were imaged with the same exposure (n=3/genotype). Compass in (A) refers to all panels: D, dorsal; V, ventral. pRPE, presumptive retinal pigment epithelium, pNR, presumptive neural retina; L, lens. Scale bar, 25 μm.

OTX2 was expressed uniformly in the pRPE and in a subset of retinal neuroblasts in E12.5 and E14.5 WT eyes (Figure 6A, C, E). By comparison, expression was nearly undetectable in *Zfp503^-/-^* pRPE at E12.5, although expression in the pNR appeared to be mildly increased (Figure 6B). In *Zfp503^-/-^* eyes, OTX2 also labeled a population of cells in the hyperplastic area of pRPE that appeared to develop into a more pNR phenotype by E14.5 (Figure 6D, F, arrows). Taken together, these observations suggest that *Zfp503* is not required for initial specification of the OC’s outer layer of neuroepithelium into a pRPE-monolayer of cells but is required for sustaining/advancing pRPE identity and the closure of the optic fissure soon after OC formation.

**Figure 6.**
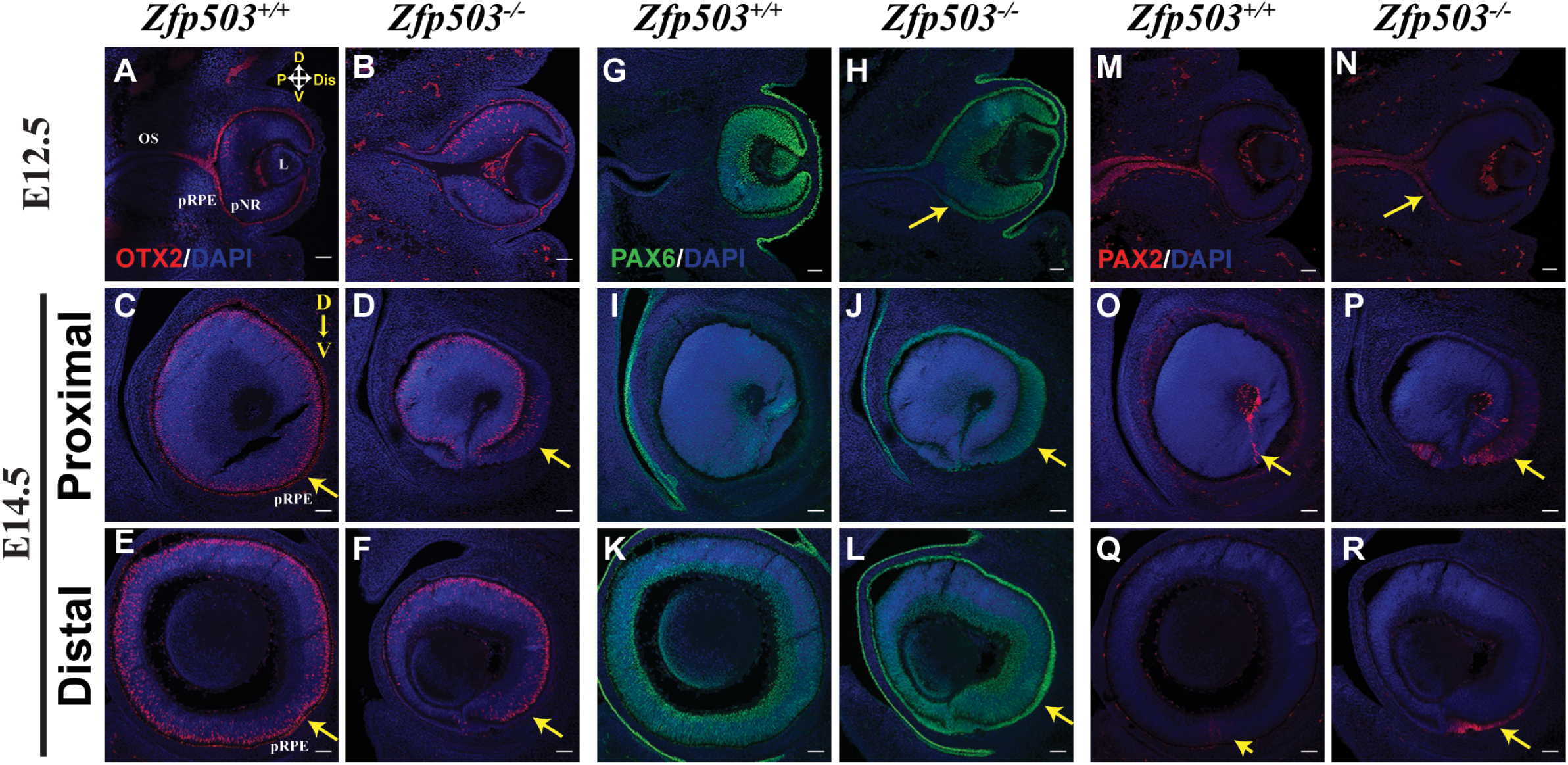
Loss of *Zfp503* expression is associated with an abnormal expression pattern of transcription factors important for RPE differentiation. Immunofluorescence for OTX2 (red, A-F), PAX6 (green, G-L) and PAX2 (red, M-R) of representative coronal sections at embryonic day (E)12.5 of WT (A, G, M) and *Zfp503^-/-^* (B, H, N) eyes, and sagittal sections at E 14.5 of WT (proximal C, I, O, and distal E, K, Q) and *Zfp503^-/-^* (proximal D, J, P, and distal F, L, R) eyes are shown. KO of *Zfp503* resulted in loss of OTX2 expression in pRPE (compare B with A). Note the scattered expression of OTX2 in the hyperplastic region (compare D, F with C, E, arrows) resembling presumptive neural retina. PAX6 expression was expanded into the optic stalk of *Zfp503^-/-^* eyes (compare H with G) and was prominently expressed in the hyperplastic pRPE (compare J, L with I, K, arrows). PAX2 expression in *Zfp503^-/-^* eyes extended beyond the optic stalk into the ventral pRPE (compare N with M, arrow) and persisted in the region of hyperplasia at E14.5 (compare P, R with O, Q, arrow). DAPI nuclear staining in blue. Confocal microscope exposure and parameters were maintained equal for comparison purposes. Compass in (A) applies to A-B, G-H and M-N, compass in (C) applies to C-F, I-L, O-R. D dorsal; V, ventral; P, proximal; Dis, distal. pRPE, presumptive retinal pigment epithelium; pNR, presumptive neural retina; OS, optic stalk; L, lens. Scale bars, 100 μm (A-B, G-H, M-N) (n=3/genotype); 200 μm (C-F, I-L, O-R) (n=2/genotype).

The homeobox transcription factor PAX6 also participates in RPE differentiation in concert with MITF and the transcription factor PAX2^33, 42^. PAX6 was detected in WT embryos in both the pNR and the pRPE (as well as the lens vesicle), particularly in the mid-to distal portions of the OC at E12.5 (Figure 6G, coronal section), consistent with previous reports. At E14.5 it was largely absent from the optic stalk and present in the pNR with less robust labeling in the pRPE (Figure 6I, K). In *Zfp503^-/-^* embryos, expression of PAX6 in pNR and lens was similar to WT at E12.5 (Figure 6H). In contrast, PAX6 expression extended well into the optic stalk from the proximal pRPE on the dorsal side of the OC (Figure 6H, arrow) and slightly more proximal in the ventral pRPE. These differences were more pronounced on sagittal sections at E14.5 (compare Figures 6J, L with I, K, arrows). Cells in the hyperplastic outer layer of the OC expressed abnormally elevated levels of PAX6, consistent with a pNR phenotype.

In contrast, PAX2 was expressed at E12.5 predominantly in the optic stalk (ventral>dorsal near the cup/stalk boundary) and qualitatively less intensely in the ventral pRPE of WT eyes (Figure 6M). However, in E12.5 *Zfp503^-/-^* embryos, PAX2 expression extended further into the pRPE of the proximal OC (ventral>dorsal) (Figure 6N, arrow). Sagittal sections of E14.5 WT eyes revealed a fine line of PAX2 immunofluorescence (Figure 6O) or a faint but detectable signal (Figure 6Q) at the, now closed, optic fissure (arrows). In *Zfp503^-/-^* embryos, PAX2 expression was more prominent and could be seen in cells of the hyperplastic pRPE (Figure 6P, R, arrows). When viewed using dual immunofluorescence labeling, the expanded expression domains of PAX2 and PAX6 could be appreciated to overlap in the proximal pRPE and dorsal optic stalk in *Zfp503^-/-^* embryos (Supplemental Figure S6), whereas they remained more distinct and exclusive at this time point in WT embryos.

Lastly, *Vax1* expression in WT embryo eyes was present in the proximal most pRPE cells within the fissure, as well as in the ventral optic stalk at E11.5 (Figure 7A, D, G, J). In *Zfp503^-/-^* eyes, *Vax1* domain was expanded into the dorsal optic stalk (data not shown) and the ventral and dorsal proximal pRPE and pNR (Figure 7M, P, S, V; data not shown). *Vax2* expression at E11.5 was strongest in the ventral retinal neuroblasts, with qualitatively less intense expression in the pRPE (Figure 7B, E, H, K) ^43^. *Vax2* was largely absent in the optic stalk. In *Zfp503^-/-^*, *Vax2* expression was similar to WT in the ventral pNR but appeared increased in the ventral pRPE and the proximal-most portion of dorsal pRPE (Figure 7N, Q, T, W, data not shown). While the expression domains of *Vax1* and *Vax2* showed some overlap in WT embryos (Figure 7C, F, I, L), this was more prominent in *Zfp503^-/-^* embryos (Figure 7O, R, U, X). These data suggest that *Zfp503^-/-^* embryos have significantly expanded domains for both these transcription factors, which partially resembles patterns of overlapping expression normally seen earlier in development.

**Figure 7.**
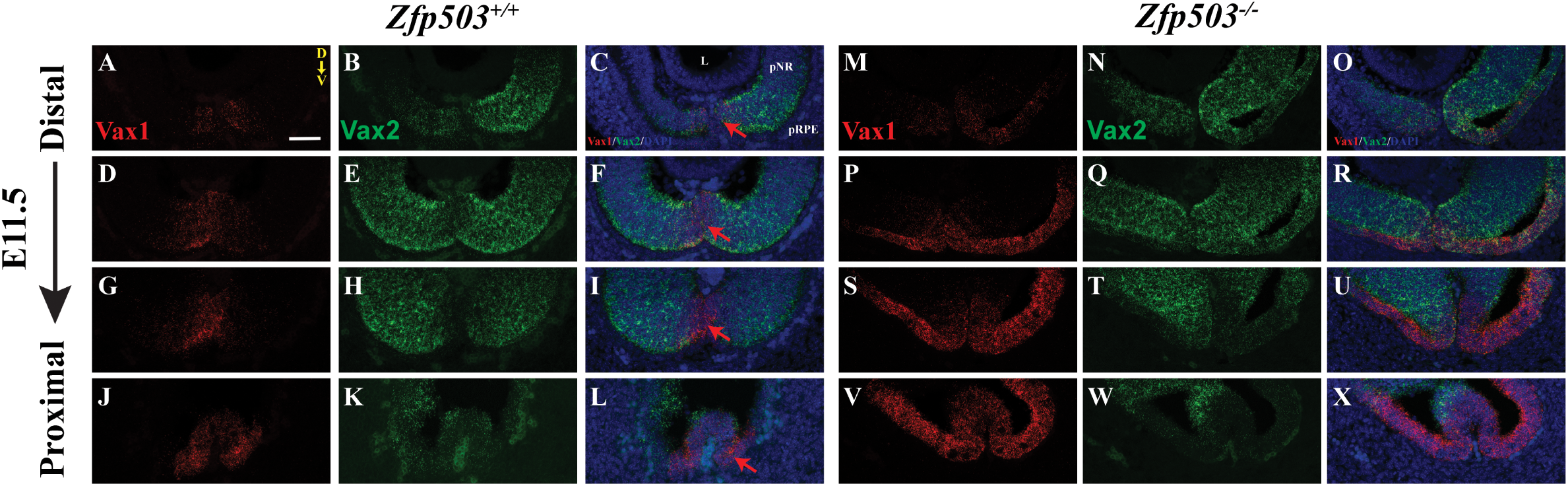
Loss of *Zfp503* is associated with abnormal *Vax1* and *Vax2* mRNA expression at E11.5. Microphotographs of representative sagittal section series, distal to proximal, of developing mouse eyes at embryonic day (E)11.5 showing *Vax1* (red, A, M, D, P, G, S, J, V) and *Vax2* (green, B, N, E, Q, H, T, K, W) expression using *in situ* hybridization. Overlay are shown in C, F, I, L, O, R, U, X, with DAPI nuclear staining in blue. In WT eyes, *Vax1* expression was limited to the very proximal ventral optic cup and the ventral optic stalk (A, D, G, J), while *Vax2* signal was mostly in the ventral retinal neuroblasts of the optic cup (B, E, H, K). In *Zfp503^-/-^* embryos, *Vax1 in situ* hybridization extended more dorsally and further into the proximal optic cup (M, P, S, V). *Vax2* expression was more prominent in ventral pRPE in addition to retinal neuroblasts and extended further into the optic stalk in KO embryos (N, Q, T, W). The images are from a single plane (n=2/genotype). Confocal microscope exposure and parameters were maintained equal for comparison purposes. Compass in (A) is applies to all panels. D, dorsal; V, ventral. pRPE, presumptive RPE; pNR, presumptive NR. Scale bar, 50 μm.

### RNA-Sequencing confirms RPE dysgenesis in *Zfp503^-/-^* mice

To better understand the pathogenesis of coloboma in our model, we performed RNA-Seq on pRPE/Choroid dissected from E11.5 WT and *Zfp503^-/-^* embryos. A total of 148 genes were significantly upregulated in *ZFP503* KO embryos compared to WT (log_2_ fold change >1); by contrast, 343 genes were significantly downregulated using the same criteria (Supplemental Tables S2 and S3). Changes in *Mitf*, *Otx2*, *Pax2*, *Pax6*, *Vax1*, *Vax2* and *Vsx2* levels (Figure 8A) were consistent with the qualitative results we observed using immunofluorescence/*in situ* hybridization and were confirmed using quantitative real-time PCR on independently dissected E11.5 pRPE (data not shown). The apparent residual expression of *Zfp503* in the KO (last panel in Figure 8A) reflected accumulation of sequencing reads in the regions flanking Exon 2 and 3. As expected, the DNA regions corresponding to the deleted Exon 2 and 3 were completely devoid of any reads (data not shown.) A volcano plot of log_10_(p value) vs. log_2_(fold change) is shown in Figure 8B. Consistent with the hypopigmentation observed in the pRPE, multiple genes directly or indirectly involved in melanin production/melanosome biogenesis were significantly downregulated (e.g., *Tyr*, *Dct*, *Slc45a2*, *Slc24a5, Gpnmb*, *Pmel*, *Gpr143, Mlana, Mlph*). Similarly, several RPE signature genes (Supplemental Table S4 ^44^) were significantly downregulated (<9.5x10^-15^, hypergeometric testing), consistent with a defect in RPE differentiation. A number of differentially expressed genes overlapped with those we previously identified by molecular profiling of optic fissure closure ^9^ (p< 3.2x10^-9^, hypergeometric testing), including *Strmn4, Insm1, Itgb8, Dcc, Tox3, Atoh7, Ascl1, Dkk3, Myb, Hes5, Fgf15, and Onecut1* (Supplemental Tables S2, S3).

**Figure 8.**
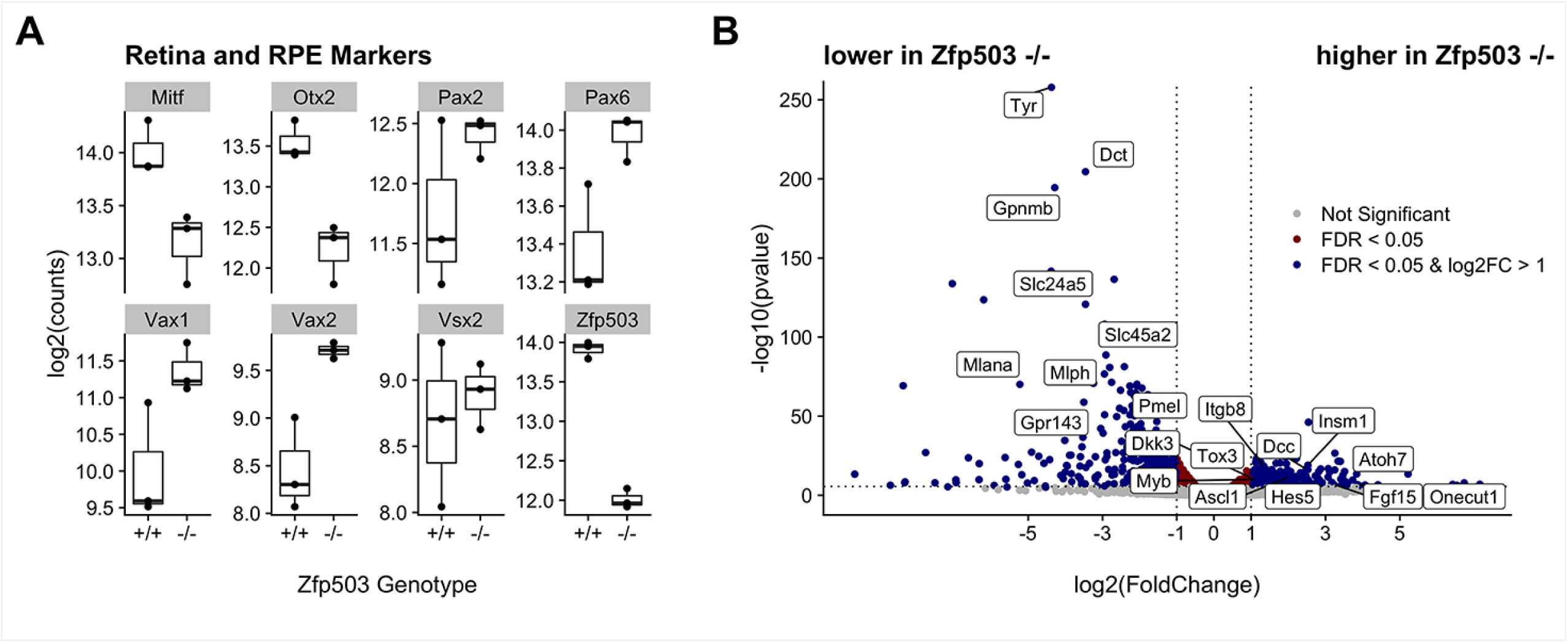
RNA-Seq from RPE at E11.5. Quantitation of several key RPE and retina mRNA transcripts in *Zfp503^-/-^* vs. *Zfp50^+/+^* embryos demonstrated changes consistent with the qualitative differences noted in immunofluorescence experiments (A). Residual expression of *Zfp503* in *Zfp503^-/-^* reflects accumulation of sequencing reads in the regions flanking the deleted Exon 2 and 3, which, in turn, were devoid of any reads (data not shown.) Volcano plot of differentially regulated genes showing changes in several key transcription factor, melanin pigmentation and genes previously identified as changing during the time of optic fissure closure (B).

### Variants in *ZFP503* and human ocular disease

To investigate whether variants in *ZFP503* were associated with congenital eye malformations in humans, we sequenced 150 affected individuals with isolated and syndromic optic fissure closure defects from 114 families. Identified variants are shown in Table 1.

**Table 1.** Heterozygous ZFP503 variants identified.

| Family | Variant<sup>1</sup> | Protein | Phenotype | Notes |
| --- | --- | --- | --- | --- |
| 98 | c.697G>C | p. Ala233Pro | Morning glory anomaly OS. Normal brain MRI/MRA, spine x-ray, kidney ultrasound, echocardiogram | Missense; rs=200297480; AF=4.9E-4; S=Tol; PP= PosD; CADD=26.1.<br>Variant also found in unaffected mother. |
| 107 | c.666C>G | p. Phe222Leu | Colobomatous microphthalmia OS;PSC OU; hemifacial microsomia; hypomelanosis of skinm scoliosis | Missense; rs747667090, AF=4.5E-0.5; S=Tol; PP=PrD; CADD=25.6. Somatic trisomy 20 identified on skin biopsy of hypomelanotic lesion.<br>Variant also found in father who was phenotypically normal by history (unavailable for exam). |
| UK | c.70G>T | p. Gly24Ter | Rod-cone degeneration, cystoid macular edema, Bockdalek hernia | Not in gnomAD. |

<sup>1</sup> Only one predicted loss-of-function heterozygous variant in an independent exome-based screen of >45,000 individuals was identified (case UK), albeit with a non-colobomatous phenotype. AF= allele frequency in gnomAD database; S = SIFT; PosD= possibly damaging; PP= PolyPhen; PrD = probably damaging; PSC=posterior subcapsular cataract; Tol=tolerated.

Heterozygous variants from families 107 and 98 were inherited from unaffected/likely unaffected parents and were present in gnomAD at low frequency in European populations (https://gnomad.broadinstitute.org; Family 107, p.Phe222Leu AF = 4.5E-5 or 1 in 36,337; Family 98, p.Ala233Pro AF = 4.9E-4 or 1 in 4082 people). These variants did not reach our lab’s threshold of evidence to be pursued further. The proband in family 107 was also found to have somatic mosaicism for trisomy 20, which may explain his unilateral colobomatous microphthalmia and hemifacial microsomia with hypopigmented skin patches. Affected individuals from these two families also did not harbor potentially pathogenic variants in genes known to cause coloboma in humans (data not shown). We also reviewed data from the United Kingdom’s 100,000 Genomes Project ^45^. Out of the cohort of approximately 45,000 individuals, one was found to have a clear loss-of-function variant (c. 706G>T, p. Gly24Ter). This 47-year-old female, however, manifested a retinal dystrophy and no evidence of coloboma. Taken together, we conclude there is unlikely to be an association between *ZFP503* coding variants and coloboma in humans.

## Discussion

The developmental processes associated with optic fissure closure have been progressively elucidated over the past half century, however, the genetics of uveal coloboma is far from well understood. Indeed, two recent studies were only able to molecularly solve approximately 8-15% of anophthalmia, microphthalmia and coloboma cases, consistent with our own laboratory’s experience ^8, 46, 47^. Higher yields for molecular testing have been noted in patients with a clearly defined syndrome ^48^. In an effort to identify candidate genes for human coloboma, we previously performed developmental profiling of optic fissure closure in mouse ^9^, a technique that others have repeated and expanded upon in mouse, chick and zebrafish in more recent years ^49–52^. Here, we show that one gene identified through our screen, *Zfp503,* when knocked out in mouse, produced a uveal coloboma associated with failure of the RPE to fully differentiate. We molecularly characterized this phenotype, showing reduced expression of critical RPE transcription factors such as MITF and OTX2 and the appearance of a neural-retina marker, VSX2, in the proximo-ventral region of the pRPE, accompanied by hyperplasia of cells resembling pNR (Figure 9).

**Figure 9.**
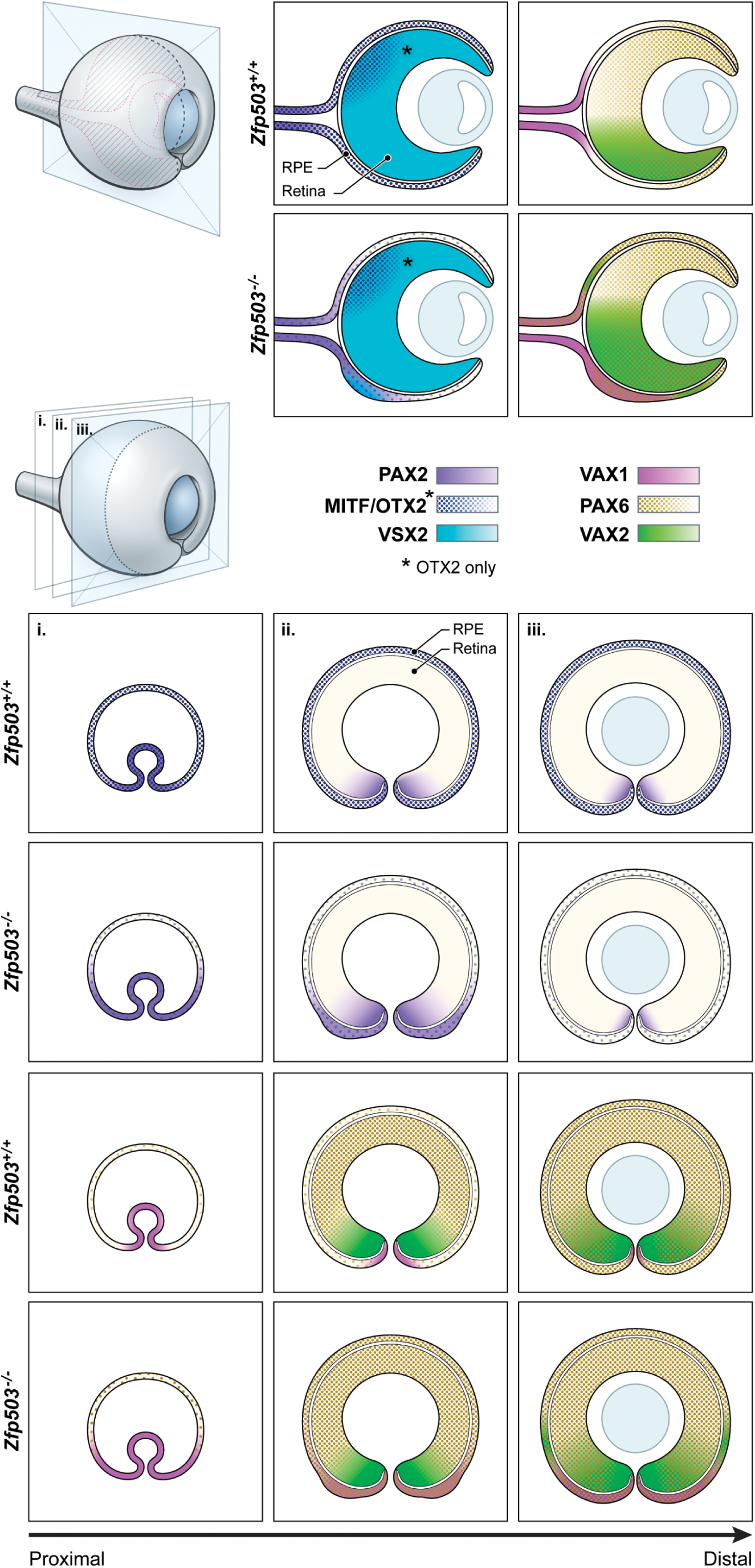
Overview of key transcription factor changes in *Zfp503^-/-^* embryos. Schematics of coronal (upper panels) and sagittal (lower panels) sections of mouse embryo eyes around the time of optic fissure closure. Three sagittal sections are shown, proximal to distal from the OS (panels i, ii, and iii, respectively). Reduction in OTX2 and MITF expression is accompanied by abnormal expansions of PAX2, PAX6, *Vax1* and *Vax2* and the adoption of a neural retina-like phenotype in the pRPE, as evidenced by VSX2 expression. Note that OTX2 and MITF changes in the presumptive RPE are very similar (and shown in one color), with the exception of possible changes in the level of OTX2 in the presumptive neural retina (denoted with an asterix).

In mouse, *Zfp503* is comprised of three exons, of which parts of exon 2 and 3 are protein coding (UCSC browser, GRCm38/mm10). The human ortholog, *ZNF503,* is composed of two exons, both of which include the protein-coding sequence (UCSC browser, GRC38/hg38). A second, noncoding *ZNF503* transcript overlaps only in the first exon. In addition, two long-noncoding RNAs (lncRNAs), *ZNF503-AS1* and *ZNF503-AS2*, each with multiple transcript variants, are present in the most recent human genome assembly. These lncRNAs are not annotated in mouse, nor have we observed them experimentally in total RNA from E11.5 mouse eyes (data not shown.) This species difference is important to note, as Chen et al. found that high levels of *ZNF503-AS1*, ergo low *ZNF503* (presumably transcript 1) expression, is correlated with increased RPE differentiation in human ^53^, the opposite of what we observed in mouse. The ocular expression pattern of ZFP503 during mouse embryogenesis is consistent with earlier studies ^9, 18, 22^ and the positive immunofluorescence signal we observed in fully differentiated mouse retina is consistent with the ganglion cell expression noted in chick by Blixt and colleagues ^16^. Although the precise cause of *Zfp503^-/-^* neonatal lethality is unclear, expression of *Zfp503* in the developing spinal cord and its role in motor neuron development could explain the atelectasis we noted in the lungs ^12^. More recently, Soleilhavoup and colleagues have demonstrated that *Zfp503* is required for proper innervation of the striatum by midbrain dopaminergic neurons in mouse and its deletion results in a nigral to pallidal lineage conversion and changes in secreted factors in the striatum ^54^; thus, we hypothesize that a developmental brain defect could contribute to perinatal lethality.

Published reports suggest that ZFP503 acts as a transcriptional repressor *in vitro*, likely through interactions with histone deacetylases and co-repressor proteins such as members of the GROUCHO family and/or ZBTB32 ^9, 12–14, 17^. The patterns of increased expression of PAX6, PAX2, *Vax2* and *Vax1* in *Zfp503^-/-^* pRPE are consistent with ZFP503 acting, either directly or indirectly, as a negative regulator of their expression (Figure 9). Notably, in breast tissue, increased ZFP503 expression results in mammary epithelial cell proliferation and more highly invasive phenotype by downregulating E-cadherin and GATA3 expression ^13, 14^.

Loss of MITF expression itself can lead to re-specification, but in the dorsal, not ventral portion of the OC ^55^. A similar phenotype mediated by an MITF- and OTX2-dependent mechanism is observed when β-catenin is conditionally knocked out in RPE, at least in the early stages of development ^56^. In the ventral pRPE, expression of the homeobox genes *Vax1* and *Vax2* is abnormally retained in the absence of MITF ^31^. Furthermore, in a triple knock-out model (*Mitf/Vax1/Vax2*) where one copy of a *Vax* gene remains, the proximal ventral pRPE transdifferentiates in a way reminiscent of *Zfp503^-/-^* embryos, implying that the combined effects of these three transcription factors are necessary for proper ventral RPE differentiation. Our finding of increased *Vax2* and *Vax1* expression in *Zfp503^-/-^* pRPE is consistent with the abnormal retention of expression seen in the MITF-KO mouse. This pattern, however, suggests that the presumed re-specification of ventral pRPE is due to another mechanism than the one observed in the three-gene KOs described by Ou *et al* ^31^.

The combined transcriptional activities of PAX2 and PAX6 are required for RPE determination ^42^. Both PAX2 and PAX6 activate the MITF-A promoter element *in vitro* and the combined absence of PAX2 and PAX6 significantly reduces MITF expression in the developing OC. The presence of expanded domains of PAX2 and PAX6 in the pRPE of *Zfp503^-/-^* E12.5 embryos suggests that, at least in the ventral-proximal region, co-expression of these transcription factors may be necessary but not sufficient for RPE determination.

RNA-Seq analysis of *Zfp503^-/-^* vs. WT RPE at E11.5 confirmed several of our phenotypic and molecular findings, including changes in genes important for early eye development (e.g., *MITF*, *OTX2*, *PAX6*, *PAX2*) and downregulation of multiple pigmentation-related genes in the KO (e.g., *Tyr*, *Dct*, *Slc45a2*, *Slc24a5, Gpnmb*, *Pmel*, *Gpr143, Mlana, Mlph*). A few differentially regulated genes are known to cause coloboma/microphthalmia/anophthalmia in humans (e.g., *Smoc1*, *Otx2, Atoh7* ^30, 57, 58^). We posit that within this gene list lie heretofore incompletely characterized genes that have a role in RPE differentiation, pigmentation and/or optic fissure closure. For example, one of the most significantly downregulated genes in *Zfp503^-/-^* is *Slc4a5*, an electrogenic sodium bicarbonate cotransporter important in brain development and cerebrospinal fluid creation ^59^. *Slc4a5* knockout mice have abnormal photoreceptor and ganglion cell numbers with retinal detachment (which may include retinoschisis) and abnormal ERGs. Importantly, although not noted in the article, *Slc4a5* mutant mice appear to have reduced RPE pigmentation in the light and electron microscopy images presented (see Figure 8 in ^59^). This observation, if confirmed, may have important implications for RPE function (e.g., transport) and for melanosome biogenesis.

What role, if any, do germline mutations in *ZNF503* have in human disease? Population-based genome databases suggest that this gene is highly constrained with a “probability of loss-of-function intolerant” (pLI) score of 0.95 and an observed/expected ratio of 0.35 (www.gnomad.broadinstitute.org), making it a good candidate for human disease. In our screen of coloboma families we found no plausible variants; in the UK 100,000 Genomes Project, we only identified one case with a clear loss-of-function allele (UK, 706G>T, p. Gly24Ter), who did not have coloboma. We note that *ZFP503* has a second ATG, in frame with the canonical translation start site, that would initiate translation with a methionine at aa 79 and produce a shorter protein isoform. Experimental data on ZFP703 that shares 53% homology with ZFP503 strongly suggest that a full-length protein and an N-terminal truncated shorter protein might be the products of *ZFP503*^19^. In that case, the variant in “UK” is predicted to allow translation of the shorter isoform from the internal ATG codon with the potential for a compensatory effect. Whether the variants reported here lead to differences in cell physiology and development awaits further study. Regardless, the percentage of inherited ocular disease caused by pathogenic mutations of *ZNF503* is (at best) very low and, given the high degree of genetic constraint and the phenotype of the KO mouse, we hypothesize that biallelic loss-of-function mutations are likely lethal.

In summary, our data support a role for the transcription factor *Zfp503* in completing the differentiation program of the RPE. In the absence of *Zfp503*, lack of RPE cells at the fissure margin results in failure to close the gap and ensuing coloboma. Based on our expression studies in the postnatal mouse eye, we hypothesize additional roles for *Zfp503* in retinal development and/or homeostasis, affecting subpopulations of cells in the RGC layer and in the INL. This hypothesis awaits further experimental testing.

## Acknowledgments

This research was supported by the intramural program at the National Eye Institute, National Institutes of Health, Bethesda, MD (Project EY000469). Dr. Panman was supported by the Medical Research Council, UK. Dr. Moshiri was supported by NIH grant K08 EY027463. Thanks goes to the NEI Histology Core for outstanding technical assistance in preparing H&E slides of mouse eyes. This research was made possible through access to the data and findings generated by the 100,000 Genomes Project. The 100,000 Genomes Project is managed by Genomics England Limited (a wholly owned company of the Department of Health and Social Care). The 100,000 Genomes Project is funded by the National Institute for Health Research and NHS England. The Wellcome Trust, Cancer Research UK and the Medical Research Council have also funded research infrastructure. The 100,000 Genomes Project uses data provided by patients and collected by the National Health Service as part of their care and support.

---

**Supplemental Figure S1. Targeted deletion of *Zfp503* in mouse results in loss of protein expression.** The entire coding region of *Zfp503* was replaced with beta-galactosidase reporter and neomycin (Neo) selection cassettes via homologous recombination in mouse embryonic stem cells (A). Proper insertion was confirmed for the left (LA) and right (RA) arms of the construct using long-range PCR (B). Hetero- or homozygous deletion of the *Zfp503* gene could be detected by PCR of tail genomic DNA based on a 216 bp WT band and a 385 bp mutant band (C). Zfp503 immunofluorescence (green) of coronal sections of WT (D) and *Zfp503^-/-^* (E) embryonic mouse eyes showed complete loss of the signal normally present in the presumptive RPE (pRPE), the optic stalk (OS) and mesenchyme (Me) at embryonic day (E)12.5. DAPI nuclear staining in blue (n=3/genotype). D, dorsal; V, ventral; P, proximal; Dis, distal. Scale bar in D, 100 μm.

**Supplemental Figure S2.** Heatmap and dendrograms showing relatedness of RNA-Seq samples is greater within a genotype (WT vs. KO) than between the two genotypes. Darker shades of blue indicate greater relatedness.

**Supplemental Figure S3. ZFP503 expression in sagittal sections of WT embryo eyes at embryonic day (E)11.5 and E12.5.** Representative microphotographs of ZPF503 fluorescence immunostaining (green, A, D) (n=3). DAPI nuclear staining in blue (B, E), along with the merged composite (C, F). Confocal microscope exposure and parameters were maintained equal for comparison purposes. Compass in (A) applies to all panels. D, dorsal; V, ventral, pRPE, presumptive retinal pigment epithelium; pNR, presumptive neural retina; OF, optic fissure; Me, mesenchyme. Scale bar, 100 μm,

**Supplemental Figure S4. Expression of ZFP503 in a subset of ganglion cell layer (GCL) and inner nuclear layer (INL) cells.** ZFP503 immunofluorescence (green) only partially overlapped with the ganglion cell marker BRN3a (red) in representative sections of the retina at postnatal day (P)41 (A-C, A’-C’, arrows). Similarly, ZFP503 labeling (green) corresponded to a subset of ISL1-labeled (red) cholinergic amacrine and ganglion cells (D-F, D’-F’, arrows) as well as of CALRETININ-labeled (red) amacrine and displaced amacrine cells (G-I, G’-I’, arrow). A’-I’ are higher magnifications of the indicated areas in A-I. DAPI nuclear staining in blue in C, C’, F, F’, I, I’. ONL, outer nuclear layer; INL, inner nuclear layer; GCL, ganglion cell layer; IPL, inner plexiform layer. Images are from single plane. Scale bar, 20 μm.

**Supplemental Figure S5. Ocular phenotype of *Zfp503^+/-^* mice.** *Zfp503*^+/-^ mice were viable, fertile and without external or obvious systemic abnormality, but exhibited excavation surrounding the optic nerve, as seen on fundoscopy (compare A with B, arrow) (A, n=4 and B, n=7) and in H&E-stained sections (compare C with D, arrow) (n=3/genotype). Scale bar, 50 μm.

**Supplemental Figure S6. Changes in the expression pattern of PAX2 and PAX6 in *Zfp503^-/-^* embryos.** PAX2 (red) and PAX6 (green) immunofluorescence in representative coronal sections of embryonic day (E)12.5 eyes from Figure 6, illustrated with both channels to demonstrate the abnormal, overlapping pattern of expression in the dorsal OS and the proximal presumptive RPE in KO embryos. Confocal microscope exposure and parameters were maintained equal for comparison purposes. Scale bar, 100 μm.

**Supplemental Table S1.** Sourcing and conditions for commercial antibodies used in this study.

**Supplemental Table S2.** Over-expressed genes overlapping with those we previously identified by molecular profiling of optic fissure closure ^9^.

**Supplemental Table S3.** Under-expressed genes overlapping with those we previously identified by molecular profiling of optic fissure closure ^9^.

**Supplemental Table S4.** RPE signature genes ^9, 44^

