## Supplementary figures and images for "Zfp503/Nlz2 is Required for RPE Differentiation and Optic Fissure Closure"

### Figure S1

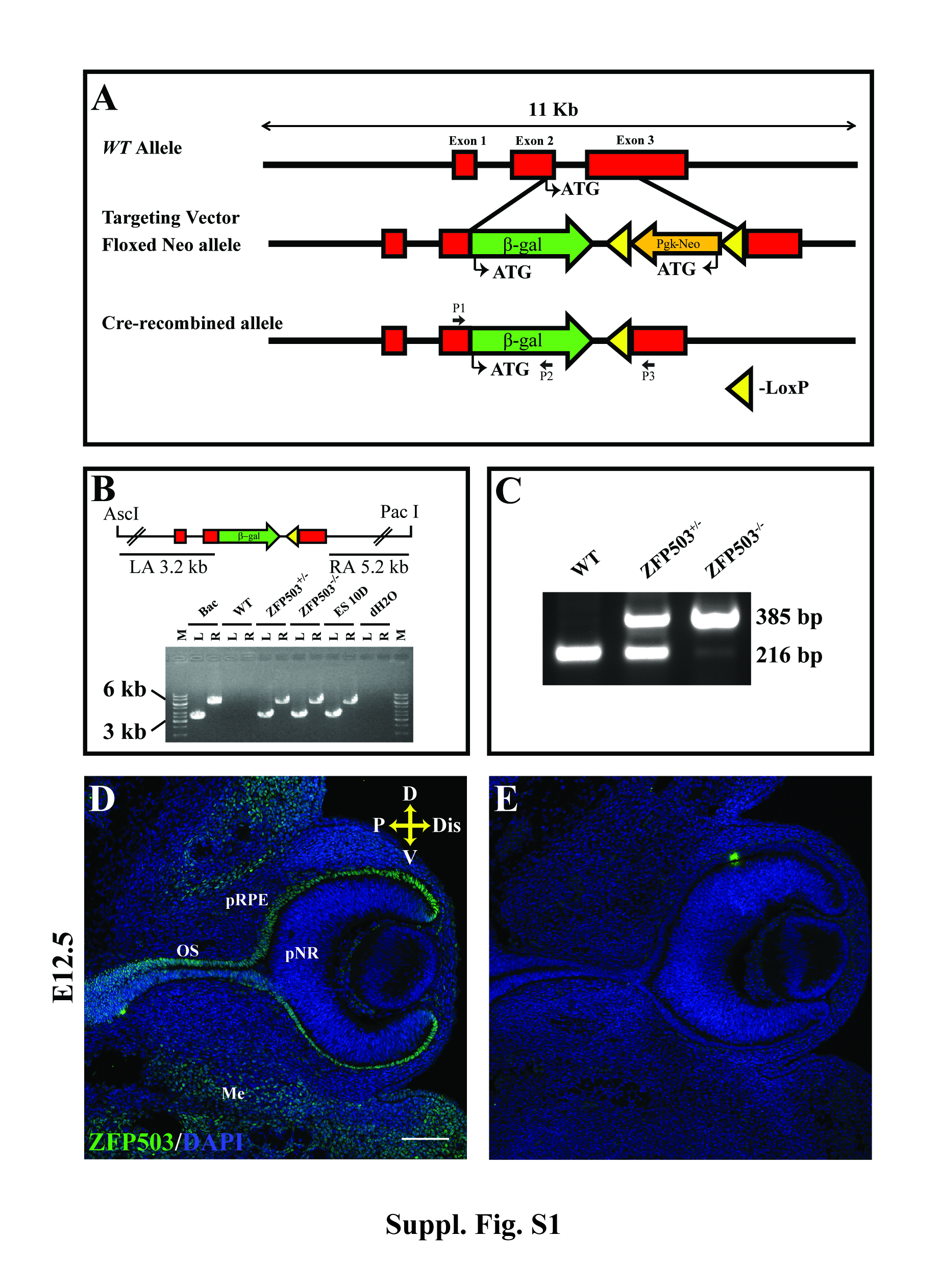

### Figure S2

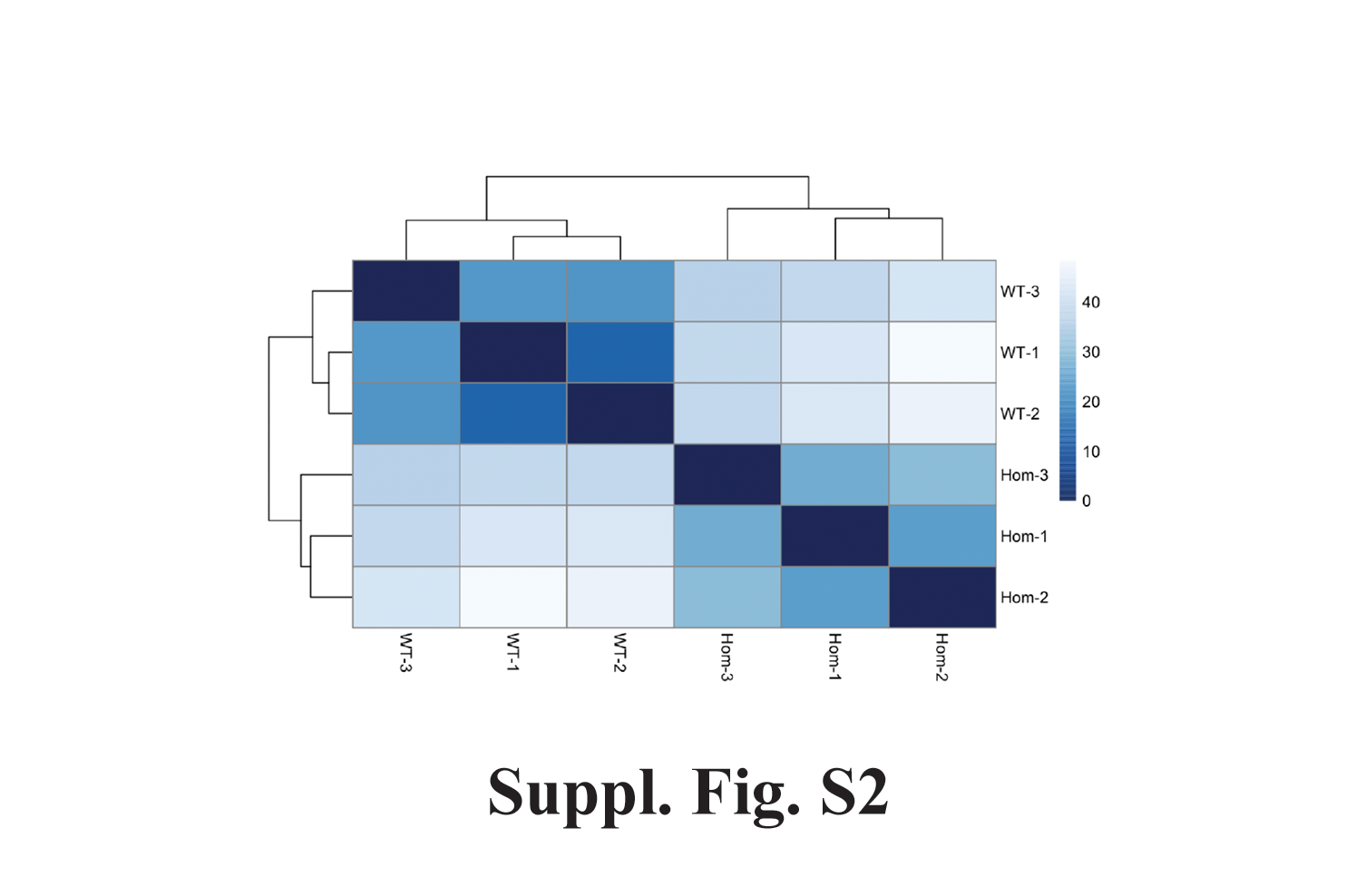

### Figure S3

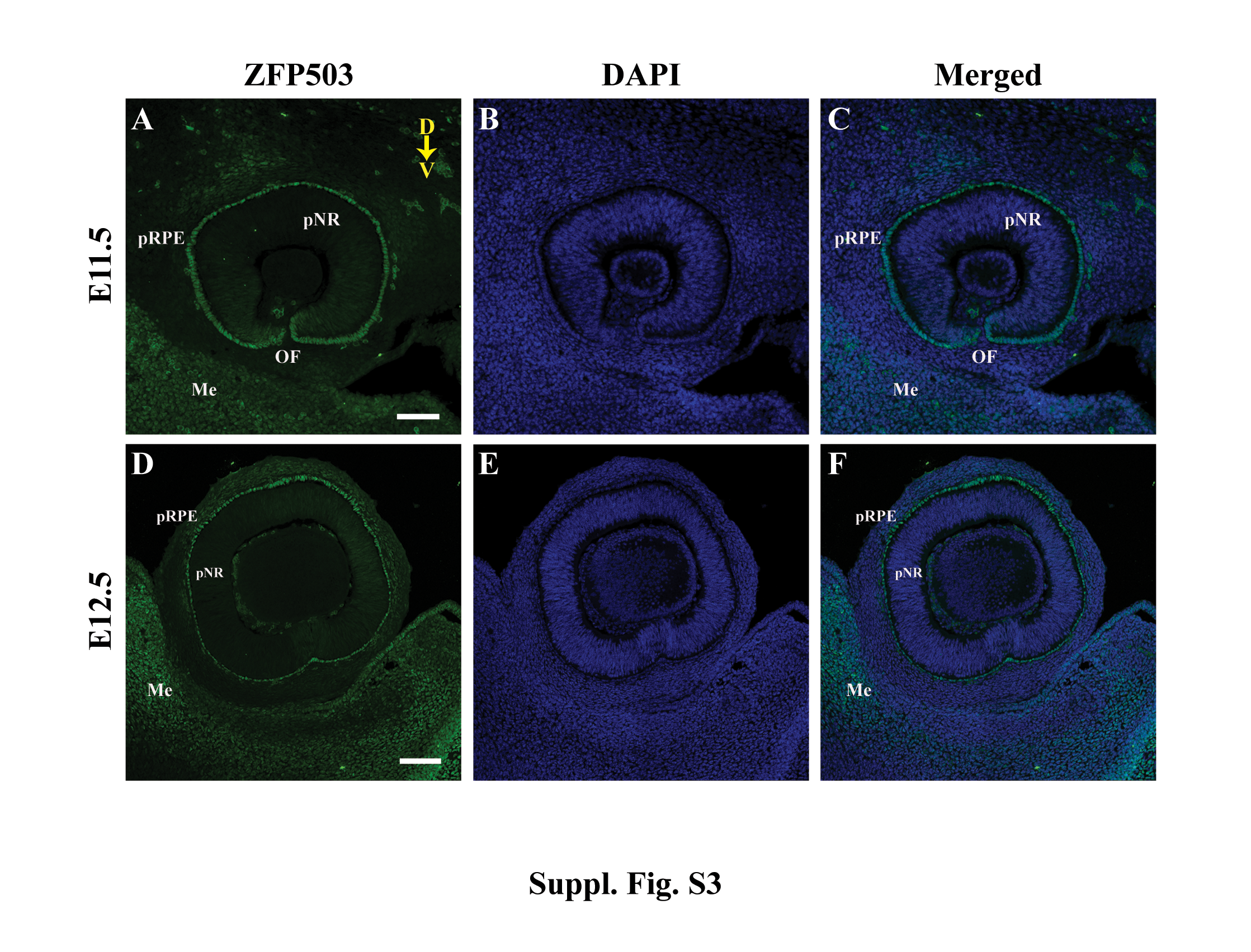

### Figure S4

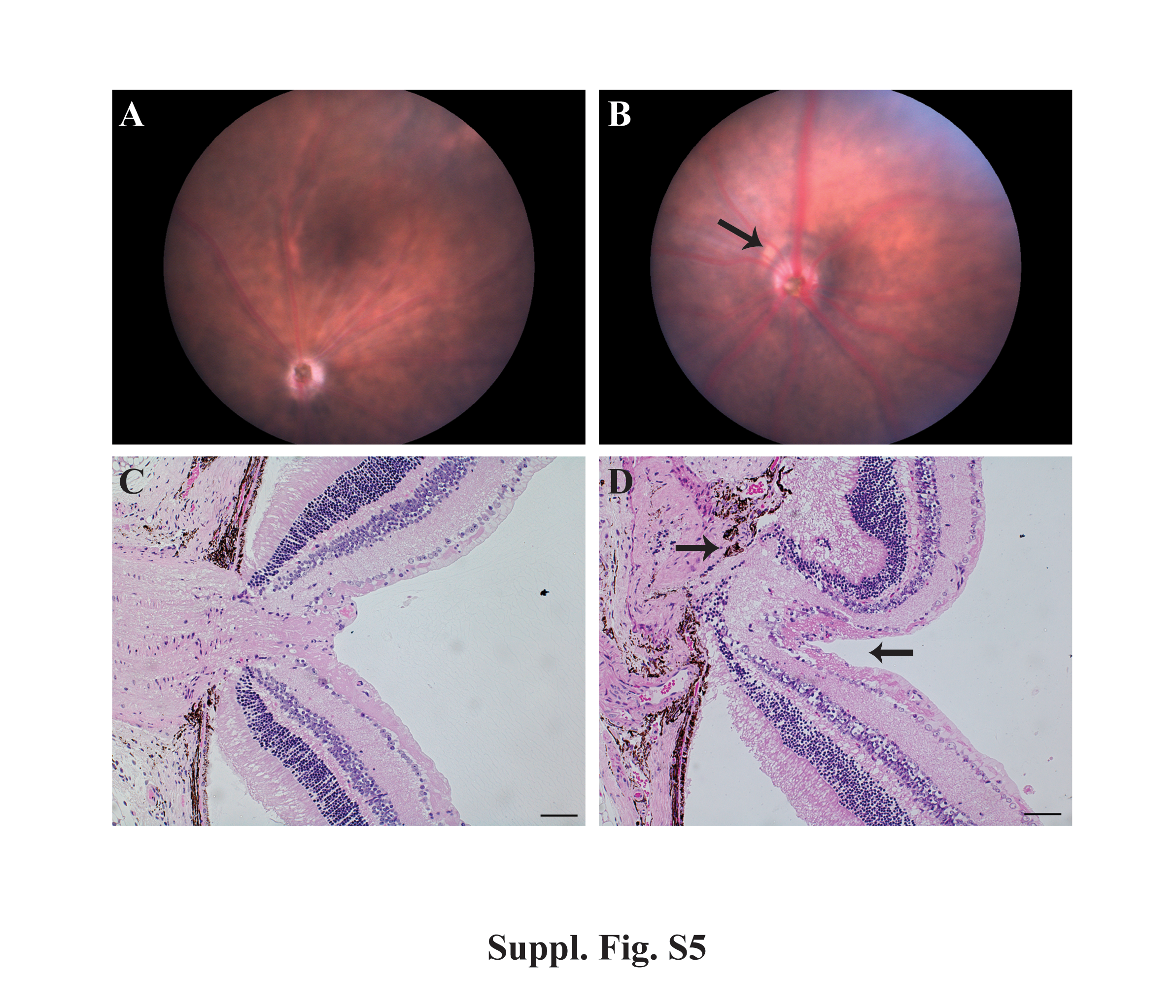

### Figure S5

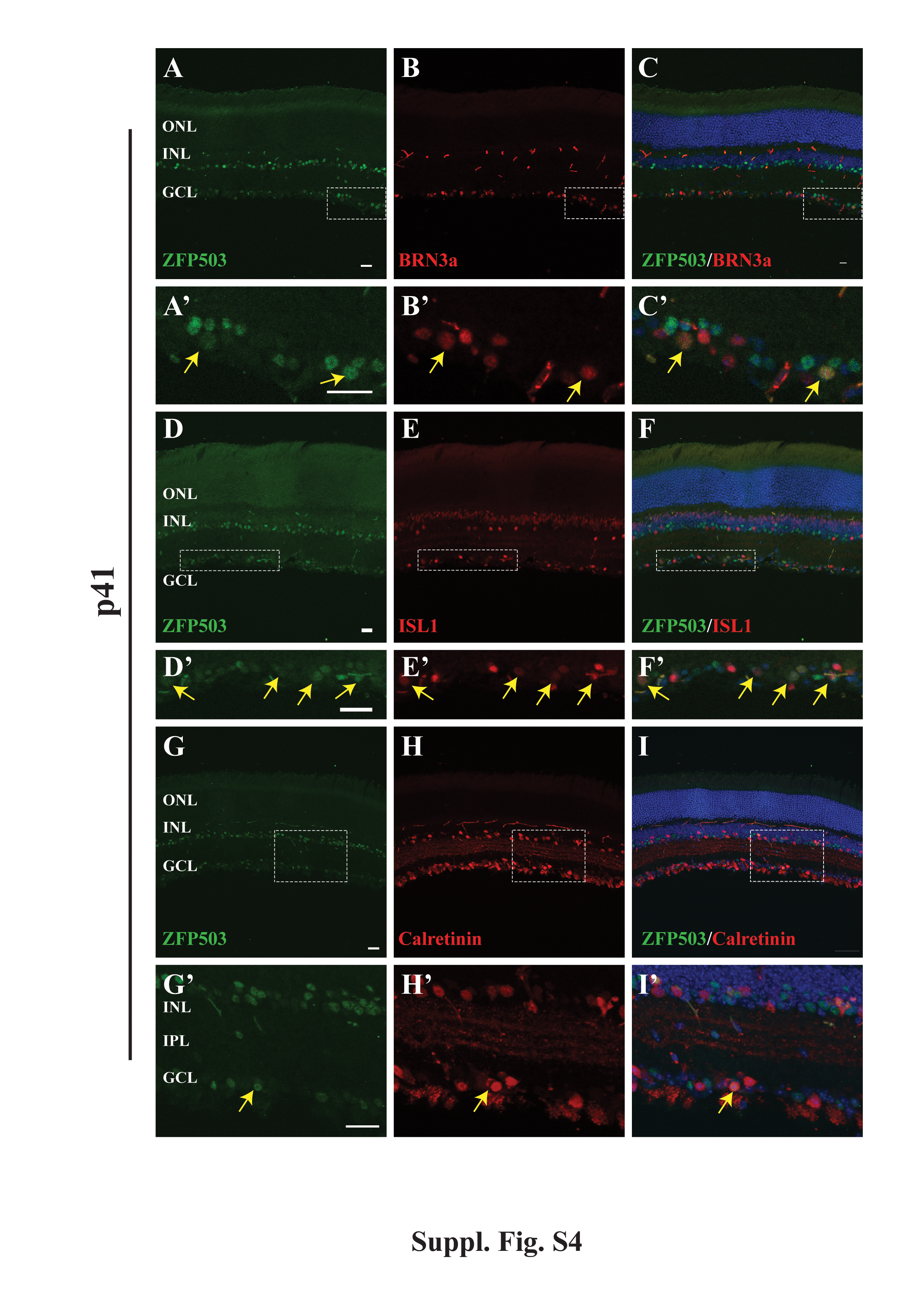

### Figure S6

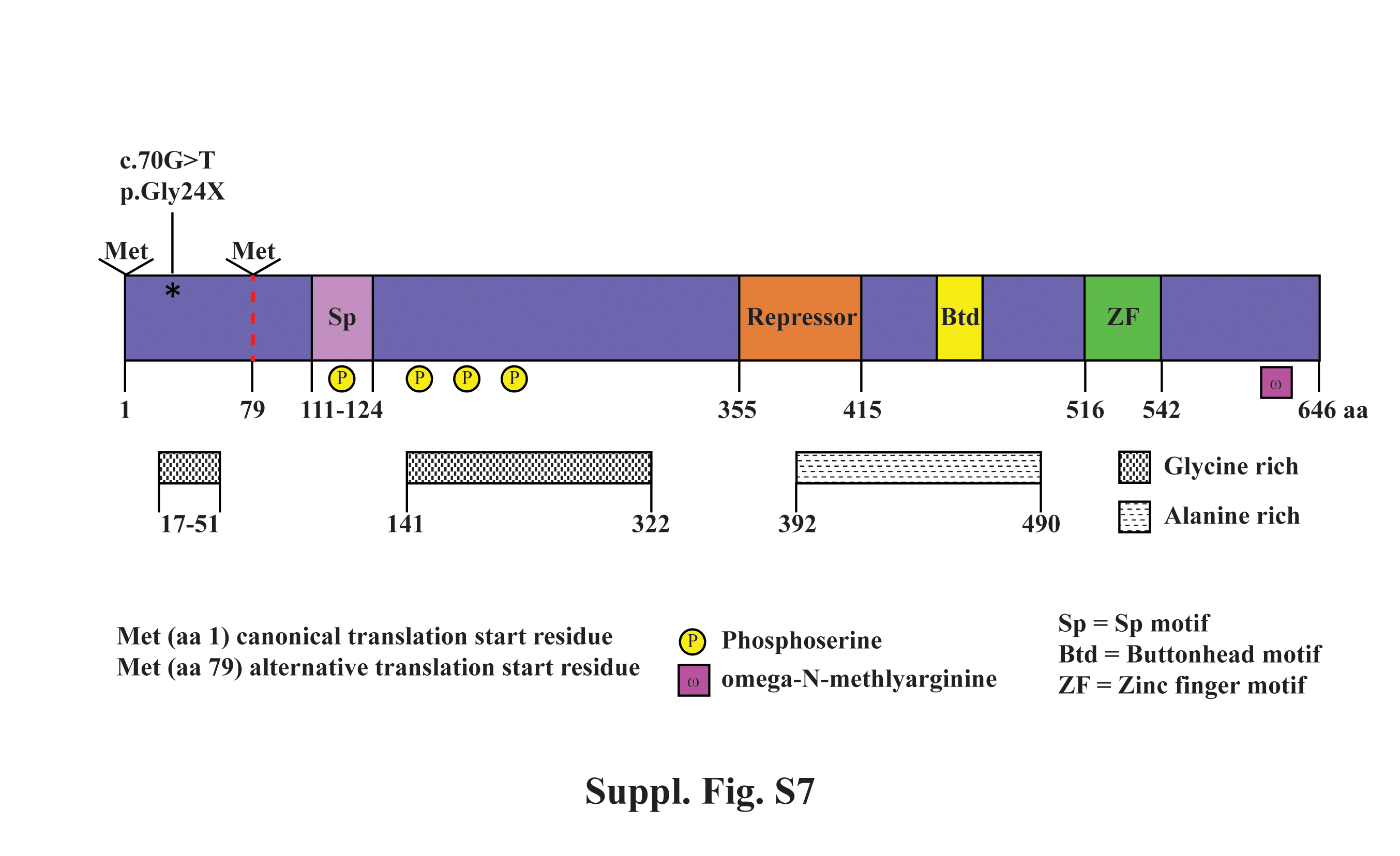

### Figure S7

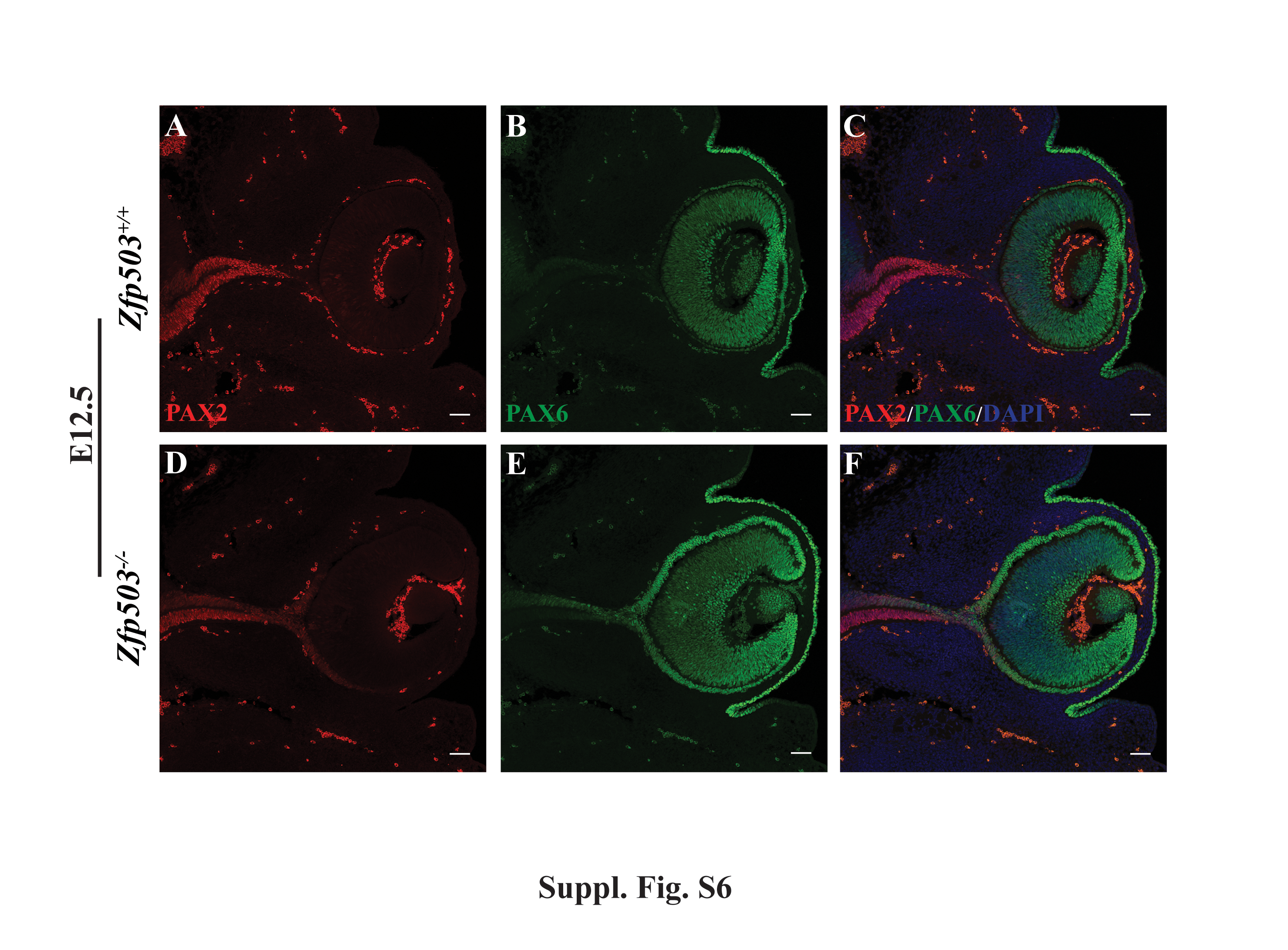
