## Supplementary Table 1 for "Zfp503/Nlz2 is Required for RPE Differentiation and Optic Fissure Closure"

**Table S1. Antibodies used in this study, with pertinent species, source and dilution information.**

| Antibody | Species | Company | Dilution |
|---|---|---|---|
| Anti-Collagen, Type IV | Rabbit | Millipore, Temecula, CA, USA | 1:1000 |
| Anti-MITF | Rabbit | Gift from Dr. Kapil Bharti (OGVFB, NEI, Bethesda, MD, USA | 1:1000 |
| Anti-OTX2 | Goat | R&D Systems, Minneapolis, MN, USA | 1:2000 |
| Anti-PAX2 | Rabbit | Invitrogen, Carlsbad, CA, USA | 1:500 |
| Anti-PAX6 | Rabbit | Covance, Princeton, NJ, USA | 1:2000 |
| Anti-CHX10/VSX2 | Goat | Santa Cruz Biotechnology, Dallas, Texas, USA | 1:2000 |
| Anti-ZNF503 | Rabbit | Sigma, St. Louis, MO, USA | 1:1000 |
| Anti-BRN3a | Mouse | Millipore, Temecula, CA, USA | 1:400 |
| Anti-CALRETININ | Mouse | Millipore, Temecula, CA, USA | 1:2000 |
| Anti-ISL1 | Goat | Neuromics, Edina, MD< USA | 1:1000 |
